# CellVADIS: Inferring cross-tissue immune cell trafficking dynamics from single-cell snapshots

**DOI:** 10.64898/2026.11.18.767220

**Authors:** Lennart Wicker, Szilard Varga, Jan Hasenauer, Frederik Graw

## Abstract

The ability of immune cells to traffic between and enter different tissues is an important hallmark for effective immune responses. However, despite advances in high-throughput single-cell profiling, identifying circulating cell populations that drive tissue-specific immunity remains elusive. Here we present CellVADIS, an optimal transport based computational framework to infer cross-tissue immune cell dynamics. By connecting single-cell measurements across tissue sites, CellVADIS enables the prediction of cell migration fates, characterization of tissue-specific trafficking signatures on circulating cells, as well as identification of tissue entry points. We benchmarked our method against synthetic data and applied CellVADIS to two cross-tissue single-cell datasets which revealed characteristic marker expression patterns that define tissue-specific trafficking signatures and ‘homing’ niches of tissue-entering CD4^+^ and CD8^+^ T cells. CellVADIS provides a flexible, open-source tool for identifying cross-tissue single-cell dynamics from various high-throughput single-cell measurements, which would support the design of vaccination and immunotherapeutic regimens to generate tissue-specific immunity.

## Introduction

Migration and motility are essential properties of immune cells, such as CD4^+^ and CD8^+^ T cells, that play an important part for the initiation and development of effective immune responses. Naïve T cells constantly circulate through lymphoid tissues in search of their cognate antigen, and, following activation and proliferation, differentiate into various T cell phenotypes, that subsequently migrate to sites of inflammation in order to mediate particular effector functions or enable long-term protection. With T cells acting locally, i.e. requiring cell-cell contacts, the targeted homing of cells to specific tissues is essential to regulate immune responses and mediate immune surveillance.

The migration and targeted homing of T cells is mediated by the expression of different cell surface proteins, such as integrins, selectins and chemokine receptors [1]. Specific receptor-ligand interactions regulate T cell homing into particular organs and tissues by mediating cell adhesion and subsequent tissue infiltration [2–5]. For example, the C-C-chemokine receptor type 7 (CCR7), which is highly expressed on naïve and central memory T cells, interacts with the corresponding ligands CCL19 and CCL21, which are highly expressed in lymphoid tissue [6]. Analogously, CXCR4 and α4β7 interact with their ligands CXCL12 and MADCAM to mediate homing to the bone marrow and gut, respectively [7,8] (Fig. **1a**). Retention of cells within particular tissues, such as lymph nodes or the spleen, is further regulated by counteracting up- and downregulation of tissue infiltration (e.g. CCR7) and egress markers, as e.g. expression of S1PR1, which binds to its ligand S1P that is highly abundant in the blood and promotes egress from lymphoid organs [9]. Interfering with chemokine receptor activity can lead to altered immune cell distribution patterns, which can result in increased sensitivity to infection, autoimmune diseases or cancer [10, 11], but could also help to resolve chronic inflammations, as e.g. in inflammatory bowel diseases [12].

**Figure 1:**
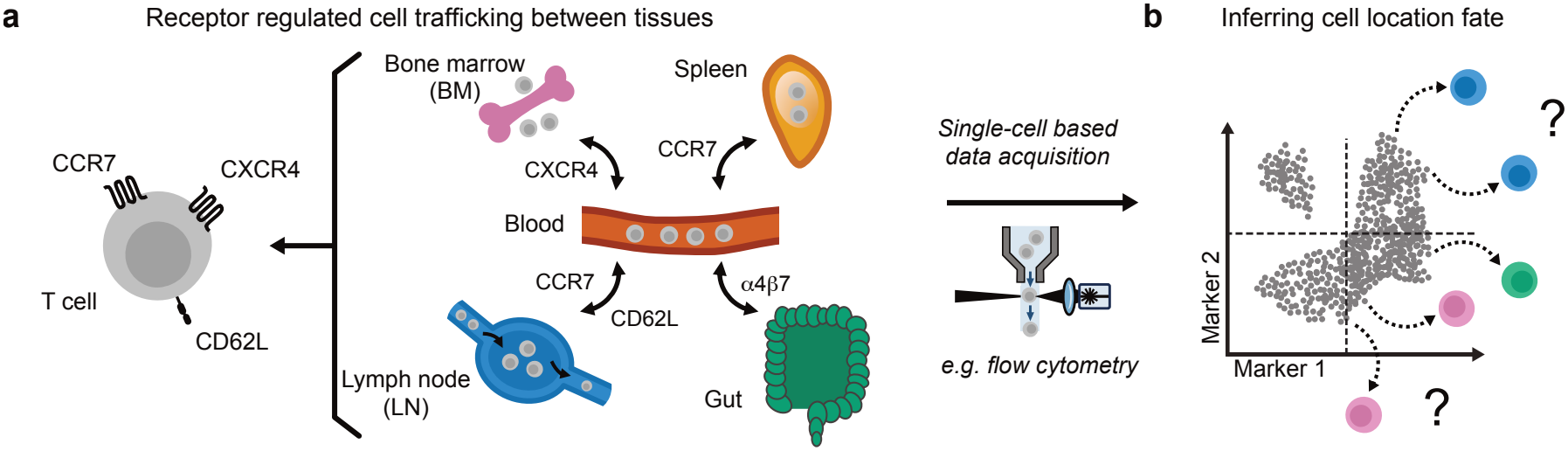
Marker-driven cell trafficking: (**a**) Dynamic expression of particular chemokine receptors and adhesion molecules on the cellular surface influences tissue-specific immune cell infiltration and circulation dynamics. (**b**) Individual marker expression levels for cells in the blood could potentially provide information to predict cell trafficking fates.

However, revealing immune cell trafficking and the establishment of tissue-specific immunity is a challenging task given that they are part of a highly orchestrated process involving the intermingled dynamics of cell proliferation, differentiation and migration [2]. T cell activation, as well as their subsequent differentiation and functionality, is strongly influenced by cell migration, as well as the localisation of cells within tissues [2, 13]. Despite experimental advances, such as photo-conversion [14] and parabiosis techniques [15], as well as *in vitro* culture systems including cellular flow [16–19], revealing cell circulation and tissue infiltration dynamics *in vivo* remains challenging. Current measurements, which rely on flow cytometry analyses or single-cell RNA sequencing, usually provide snapshot information of cell populations for different tissues at a single time point, lacking the ability to obtain continuous time-resolved measurements of cell circulation dynamics. In addition, relying on pre-defined cell populations, as for example obtained by boolean-gating strategies for flow cytometry data, would ignore the continuous receptor expression levels present in the data, which might contain information about individual cell fates [20].

Several computational methods, such as Slingshot [21] and Wishbone [22], have been developed to infer cell-state trajectories from high-dimensional single-cell measurements, particularly scRNA-seq data [23–25]. These trajectory inference methods exploit gradual changes in transcriptomic profiles to establish relationships between cells and thereby reconstruct putative trajectories through cell-state space. However, most methods are limited to detecting cell trajectories within a single snapshot, which could in theory be extended to combine samples by pooling cells [26], but do not directly integrate tissue- or temporal information to infer dynamics across samples.

Optimal transport (OT), a mathematical theory that compares probability distributions in order to find the most efficient mapping between them, has emerged as a powerful tool for connecting such high-dimensional cellular measurements. Methods, such as WaddingtonOT [27] or moscot [28], have helped to reveal time courses of cell developmental pathways by connecting time-resolved scRNAseq-measurements, with the latter enabling the integration of multi-omics data. However, these methods primarily infer sequential coupling between cell states across temporally ordered samples. Moreover, they generally describe directional progression through cell-state space, whereas immune-cell trafficking can involve competing transitions to multiple tissues. This fundamental difference limits their applicability to reconstruct cross-tissue trafficking dynamics.

In order to overcome these limitations, we developed novel methods based on optimal transport theory to allow the inference of cross-tissue T-cell dynamics. The approach enables the prediction of cell trafficking fates based on individual cellular expression profiles, as obtained by flow cytometry or scRNAseq-measurements (Fig. **1**b). After validating and benchmarking our method based on synthetic data by developing a stochastic, individual cell-based mathematical model describing cell differentiation and migration regulated by the continuous, individual expression of cell-based surface markers, we applied the approach to two published cross-tissue single-cell datasets relying on mass cytometry and scRNAseq, respectively [29, 30]. Our method enables the prediction of T cell trafficking fates, as well as the identification of essential markers and cell populations within individual tissues that help to infer tissue-specific differentiation pathways.

## Results

### Inferring cell trafficking dynamics by optimal transport

To determine the relationship between individual cells across different tissue sites, we based CellVADIS on optimal transport (OT), which is a mathematical approach that links two probability distributions by finding the most efficient way to transport mass between them. Applied to immune-cell trafficking, OT can link distributions of cellular expression profiles measured in different tissues. With the trafficking of immune cells between different tissues guided by physiological trafficking routes and not all tissues being directly connected, the blood serves as a central transit compartment through which immune cells can reach different lymphatic and peripheral tissues. This organization gives rise to a multi-target transport problem, in which cells in the blood can transit to any of *L* different target tissues (e.g., LN, gut, and spleen; Fig.2a). Inferring the tissue-specific transition potential of circulating cells therefore requires relating the cellular expression profiles in the blood to the distributions observed in all target tissues.

Based on unbalanced OT [31], which accounts for the fact that not all blood cells can be mapped to all target tissues due to unobserved tissues or cell proliferation, we developed two approaches to infer tissue-specific trafficking patterns from individual cellular expression profiles: *StitchOT* solves a separate OT-problem between blood cells and each individual target tissue with the resulting OT-maps subsequently combined using tissue-specific weights to obtain a global OT-map (Fig. 2b). *StitchOT* works from the perspective of a blood cell by estimating blood cell specific affinities (so called marginals) towards target tissues, while relaxing the constraint that each blood cell needs to be mapped to a specific tissue. By normalizing the computed OT-maps we obtain a fate matrix *F*, which assigns each blood cell a probability to migrate to a specific target tissue. This allows us to obtain tissue-specific transition probabilities based on cell expression profiles, which enable the identification of trafficking signatures on blood cells, corresponding to ‘postal codes’, that define the potential traffic direction of cells. In contrast, *JointOT* pools cells from all target tissues and solves a single OT-problem, allowing blood cells to be jointly matched across competing tissue destinations (Fig. 2c). In comparison to *StitchOT, JointOT* works from the perspective of the target cell by assigning each target cell an affinity (marginal value) towards blood cells. This allows the identification of ‘homing niches’ inside specific tissues that characterize potentially recently immigrated or constantly circulating cells due to their higher similarity to blood cells.

**Figure 2:**
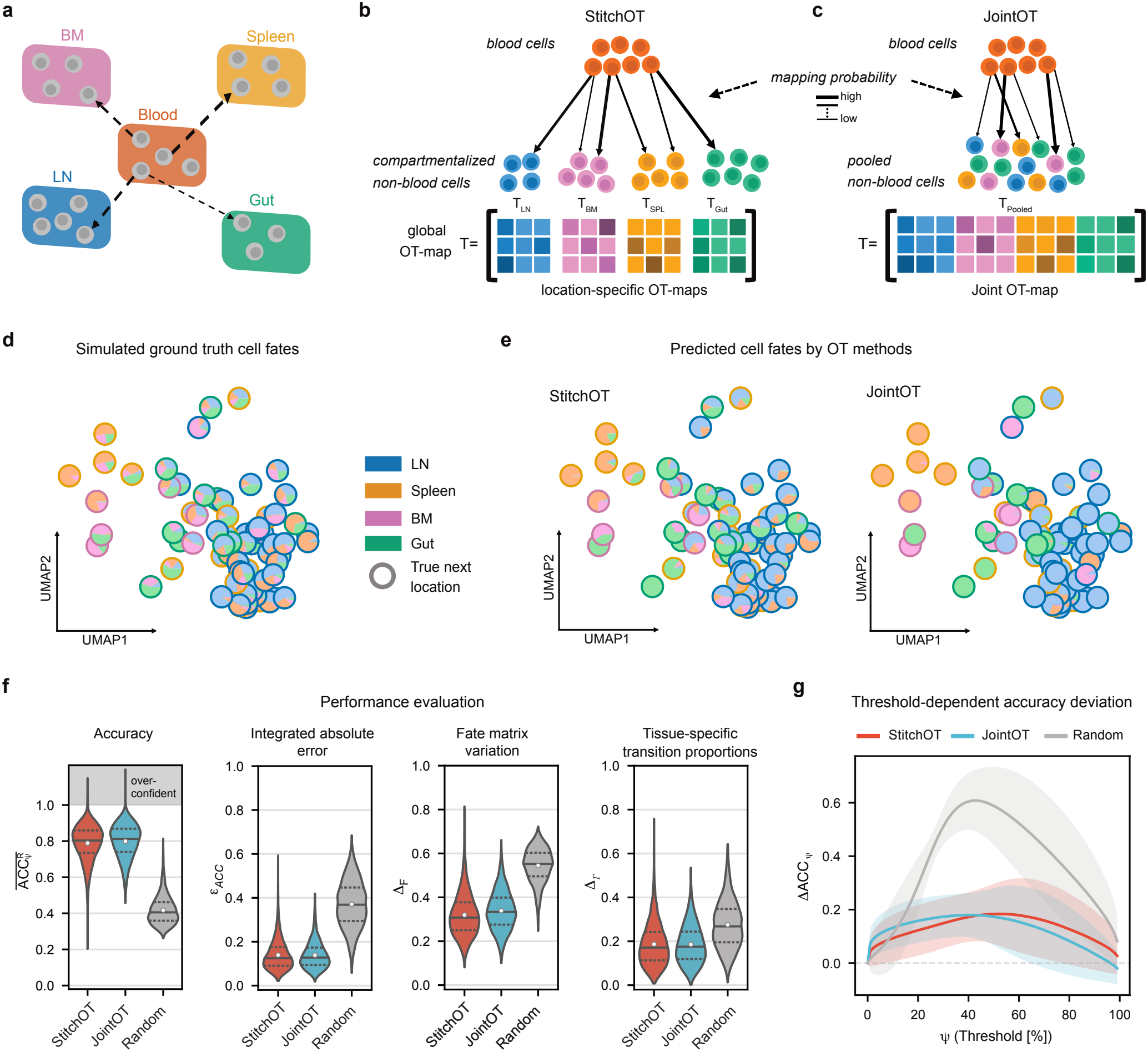
Extended optimal transport methods to infer cell trafficking fates: (**a**) Predicting trafficking fates of blood cells to sampled tissue sites by linking individual cells using extended OT methods. (**b-c**) Sketch of the concept of *StitchOT* (**b**), relying on the generation of a global OT-map by concatenating individually determined location-specific OT-maps, as well as *JointOT* (**c**), which generates a combined OT-map based on pooled tissue cells. (**d-g**) Method evaluation showing simulated ground truth transition probabilities (**d**) and predicted cell fates (**e**) using either *StitchOT* or *JointOT* for a single example comprising 100 randomly sampled blood cells at 24h post simulation start. Each pie chart represents a single blood cell, with the different colors indicating the true (**d**) and predicted (**e**) trafficking probability for the specific tissues, and the coloured circle denoting the true next location the cell would migrate to within the simulation. (**f**) Systematic method evaluation over 10^4^ simulations comparing *StitchOT* and *JointOT* to randomly shuffled OT-maps showing different performance measures including the relative average threshold accuracy, 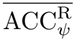, and integrated absolute error, ε_ACC_, as well as the total variation in transition probabilities, Δ_F_, and tissue-specific transition proportions, Δ_r_, respectively (see also Suppl. Fig. S2). (**g**) Mean paired differences (and standard deviation) between the ground truth 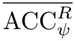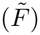 and the inferred ACC_ψ_ threshold accuracies over all possible thresholds *ψ ∈* (0, 1). Values below 0 indicate method overconfidence. (Parameters for simulation see Suppl. Table S2,S3,S4).

In summary, by their different definitions, *StitchOT* and *JointOT* either enable the identification of tissue-specific trafficking signatures in the expression profiles of blood cells to guide cells towards specific tissues, or to characterize entry points and circulation potential of tissue cells.

### CellVADIS accurately predicts cell trafficking fates

To evaluate the ability of *StitchOT* and *JointOT* to predict cell trafficking fates, we first considered synthetic data generated by a newly developed stochastic, individual cell-based mathematical model. In this model, cellular trafficking and differentiation dynamics are regulated and characterized by the continuous expression of individual cell-surface markers and chemokine receptors (Suppl. Information Text S1, Suppl. Fig. S1). The circulation dynamics of 10^4^ *−* 10^5^ cells was simulated over 30 hours, neglecting cellular turnover and maintaining a constant number of cells throughout the time course of the simulation. At 24 hours, samples were taken from all simulated tissues and the ability of our developed OT-methods to predict the subsequent tissue a sampled blood cell would migrate to was assessed based on various measures (see *Materials and Methods*).

For each simulated blood cell, the tissue-specific transition probabilities are calculated based on the individual marker expression levels (Fig. 2d), and compared to the predictions performed by the developed OT-methods (Fig. 2e), as well as the true next location a cell would migrate to. Both OT-methods, i.e. *StitchOT* and *JointOT*, show comparable levels of relative accuracy, 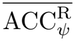, in predicting the true next location of a blood cell, reaching levels of 0.788 (0.604-0.922) (mean and *q*_0.05_-*q*_0.95_-quantile range) and 0.799 (0.626-0.931), respectively, across 10^4^ independent simulations. Thereby, both OT-methods perform ~ 2-fold better than random predictions by randomly shuffled OT-maps (Fig. 2f) with ACC 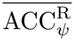-levels of 0.417 (0.261-0.813). While inferred cell-based transition probabilities,. *Δ*._F_, deviate absolutely on average by 31.9 percentage points (pp) (*StitchOT*) and 33.9 pp (*JointOT*), the predicted fraction of cells transitioning to a particular tissue,. *Δ*._r_, only show slight deviations from the ‘true’ transition proportions by on average 18.6%pt and 18.5%pt, with both OT-methods showing similar performance and clearly outperforming random assignments (Fig. 2f). Furthermore, the integrated absolute error between the practically possible optimal accuracy and the method based accuracy, ε _ACC_, indicated equal performance between both methods (Fig. 2f). However, with the predicted tissue-specific transition probability assumed to be accurate if it is larger than a defined threshold ψ, *JointOT* indicated to be on average slightly overconfident for values & *ψ ≳* 0.5 compared to *StitchOT*, with reversed roles for both methods given ψ *≲* 0.5 (Fig. 2g).

Overall, despite these minor differences, both methods enable us to robustly predict the trafficking fate of individual blood cells based on individual cellular expression profiles. Additional sensitivity analyses, which evaluate the impact of method variations on their individual performance, confirmed the robustness of the chosen approaches (Suppl. Fig. S2).

### Impact of unobserved tissue sites on cell fate prediction

Experimentally sampled tissue will usually only represent a fraction of all tissues to which blood cells can migrate. To assess how unobserved tissue sites affect cell fate predictions by *StitchOT* and *JointOT*, we considered incomplete sampling of tissues accessible to blood cells. Unobserved tissue sites can affect inference in two ways: (i) cells destined for unobserved tissues may be incorrectly assigned to sampled tissues, and (ii) these assignments may also alter predictions for cells migrating to sampled tissues (Fig. 3a,b). The impact is expected to depend on whether migration to unobserved tissues is associated with distinctive marker expression profiles. If migration to unobserved tissue sites is not controlled by specific expression profiles, or controlled by expression profiles as they are characteristic for sampled tissues (i.e. representing a ‘twin tissue’ as e.g. unobserved LN), cell fate probabilities will still be correctly assigned by our OT methods, except for overestimating the tissue-specific transition proportion for the sampled ‘twin tissue’ in the latter case. However, if cells within the unobserved tissue are characterized by marker profiles distinct from all other cells (Fig. 3c), methods that are based on distances in marker space, such as optimal transport, will by default lead to potential mis-mapping of cells, as they will assign cells to the least wrong destination. In order to assess the extent to which cell fate predictions are affected in such a scenario, we extended our simulation framework, in which migration to an unobserved tissue *U* is driven by a unique marker Λ that at the same time represents an antagonist for migration of blood cells to any of the other tissues measured, with the grade of separation regulated by parameter *γ* (see *Materials and Methods*). This results in disparate marker profiles between sampled and ‘unobserved’ tissues driven entirely by Λ that is highly expressed within the unobserved tissue site *U* (Fig. 3c, Suppl. Fig. S3). The corresponding fate matrix *F*, will then be extended to additionally account for tissue *U*. Method assessment is therefore based on the full fate matrix, as well as the conditional fate matrices for blood cells not trafficking to *U* (see *Materials and Methods* for details).

**Figure 3:**
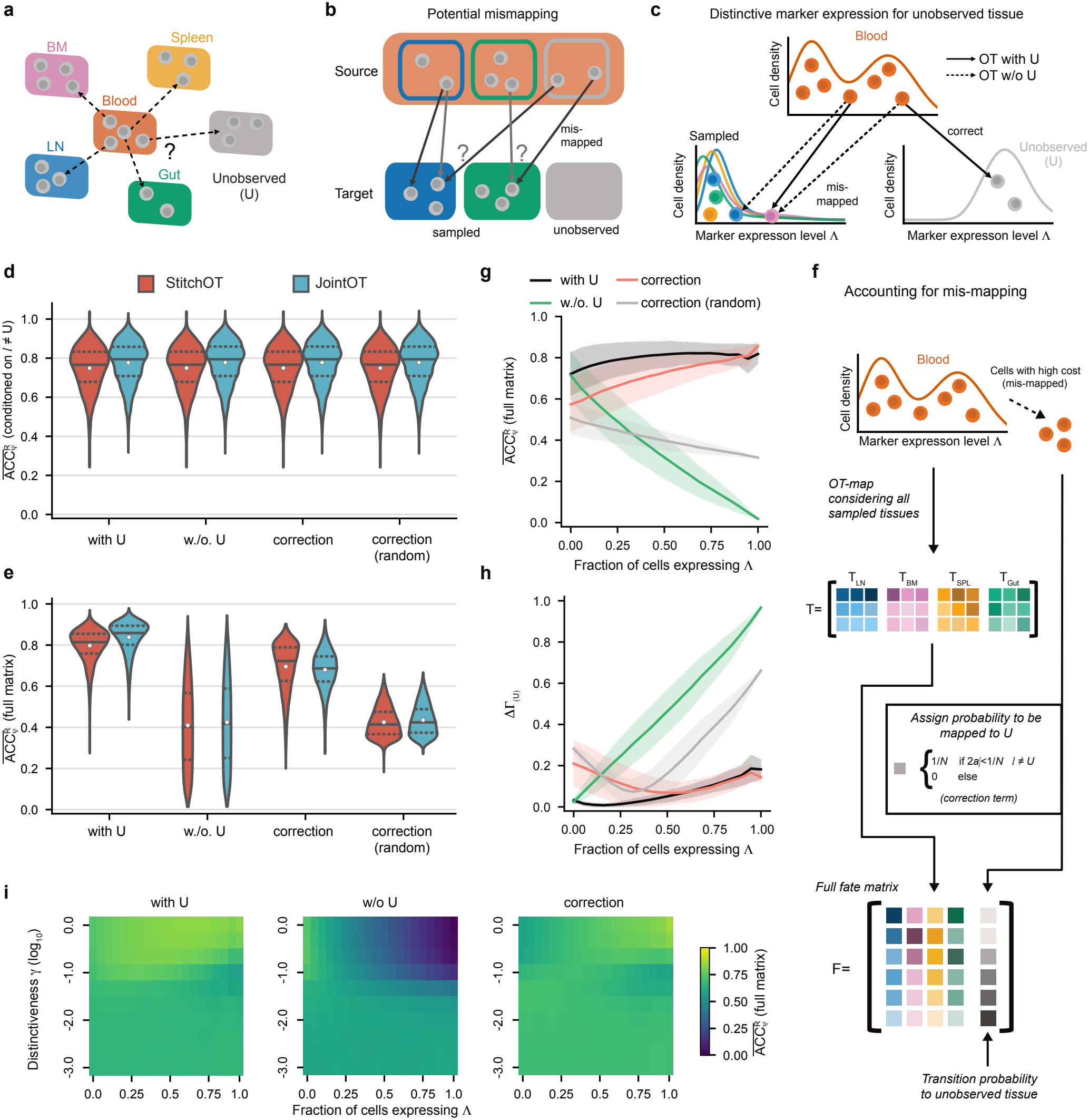
Cell trafficking fate prediction given unobserved tissues: (**a-b**) Incomplete sampling of tissue sites (**a**) can influence trafficking fate predictions due to mis-mapping of cells migrating from the blood to the unobserved tissues, and interfering with the mapping of cells to sampled tissue sites (**b**). (**c**) Presence of unique markers specific for the unobserved tissue site influences the mapping method. (**d-e**) Average threshold accuracy, 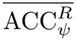, based on the conditional fate matrices, i.e. only considering cells not trafficking to *U* (**d**) and the full fate matrix considering all blood cells (**e**) for *StitchOT* and *JointOT* given different approaches. (**f**) Sketch of extending the fate matrix *F* by the probability of a cell to be mis-mapped. (**g**) Average threshold accuracy dependent on the fraction of cells expressing the unique marker Λ characterizing transition to the unobserved tissue site *U* for the different conditions for *StitchOT*. (**h**) Corresponding deviation of the predicted to the true fraction of cells transitioning to the unobserved tissue, i.e, defined by the tissue-specific transition proportion to *U* given by. Δ_Γ(*U*)_ = |Γ(*U*) *-* Γ^OT^(*U*)|. (**i**) Average threshold accuracy for different levels of the fraction of cells transitioning to the unobserved tissue site *U* and the marker distinctiveness, i.e. ϒ. For each combination 1000 simulations were performed. (**d-e**,**g-h**: *n* = 20000 simulations with a sharp distinction for Λ-mediated migration (ϒ = 1), **g-h**: solid line=mean, shaded area=standard deviation).

Applying our OT-methods to the simulated data with and without the unobserved tissue site, we found that cell fate predictions for those cells that would actually migrate to sampled tissues are not affected by the missing tissue, irrespective of the method used (Fig. 3d), which is expected based on theoretical considerations (see Suppl. Information Text S2). However, across all blood cells, substantial mis-mapping occurs, with cell fate prediction heavily impaired with the accuracy halved for both OT-methods down to 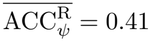 (Fig. 3e, Suppl. Fig. S3).

To account for this mis-mapping, we extended our OT-methods by the probability of a cell to be mis-mapped, i.e., exploiting the fact that optimal transport will characterize mis-mapped cells by a high cost associated with a mapping to any of the sampled tissues (Fig. 3f, *Materials and Methods*). We found that the accuracy of cell fate prediction can be restored, with slightly better performance for *StitchOT* (0.70; 0.46-0.84) (mean; *q*_0.05_-*q*_0.95_-quantile range)) compared to *JointOT* (0.68; 0.51-0.82). Hereby, the considered adaptations perform substantially better than randomly sampled mis-mapping probabilities (Fig. 3e). This means that the adapted methods are able to robustly predict the fraction of blood cells that would not be trafficking to any of the sampled tissue sites, significantly restoring the prediction accuracy to levels as if the unobserved tissue site was measured, if more than 20% of the blood cells would actually migrate to unobserved tissue sites (Fig. 3g,h). The restoration of accuracy in cell fate prediction is thereby also independent on the distinctness of the disparate marker profiles that distinguish cells from the unobserved and sampled tissue sites (Fig. 3i).

### CellVADIS identifies homing niches of blood cells within particular tissues

Given the successful benchmarking based on synthetic data, we used CellVADIS to determine tissue-specific trafficking patterns in circulating T cells by applying it to a published mass cytometry (MASS) dataset profiling T cells across human tissues [29]. Considering measurements of peripheral blood mononuclear cells (PBMC) as the source tissue, we used CellVADIS to predict transition fates towards lymphoid and non-lymphoid tissues, including the tonsils, lung, skin, colon, spleen and liver (Fig. 4a), accounting for differentiation, activation and trafficking markers. Cell fate predictions for 10^4^ CD8^+^ T cells (Fig. 4b) indicate that ~ 40% of the cells have a high affinity towards the liver, followed by the spleen (25%) and tonsils (18%) (Fig. 4c), with similar observations made for CD4^+^ T cells (Suppl. Fig. S4a,b).

**Figure 4:**
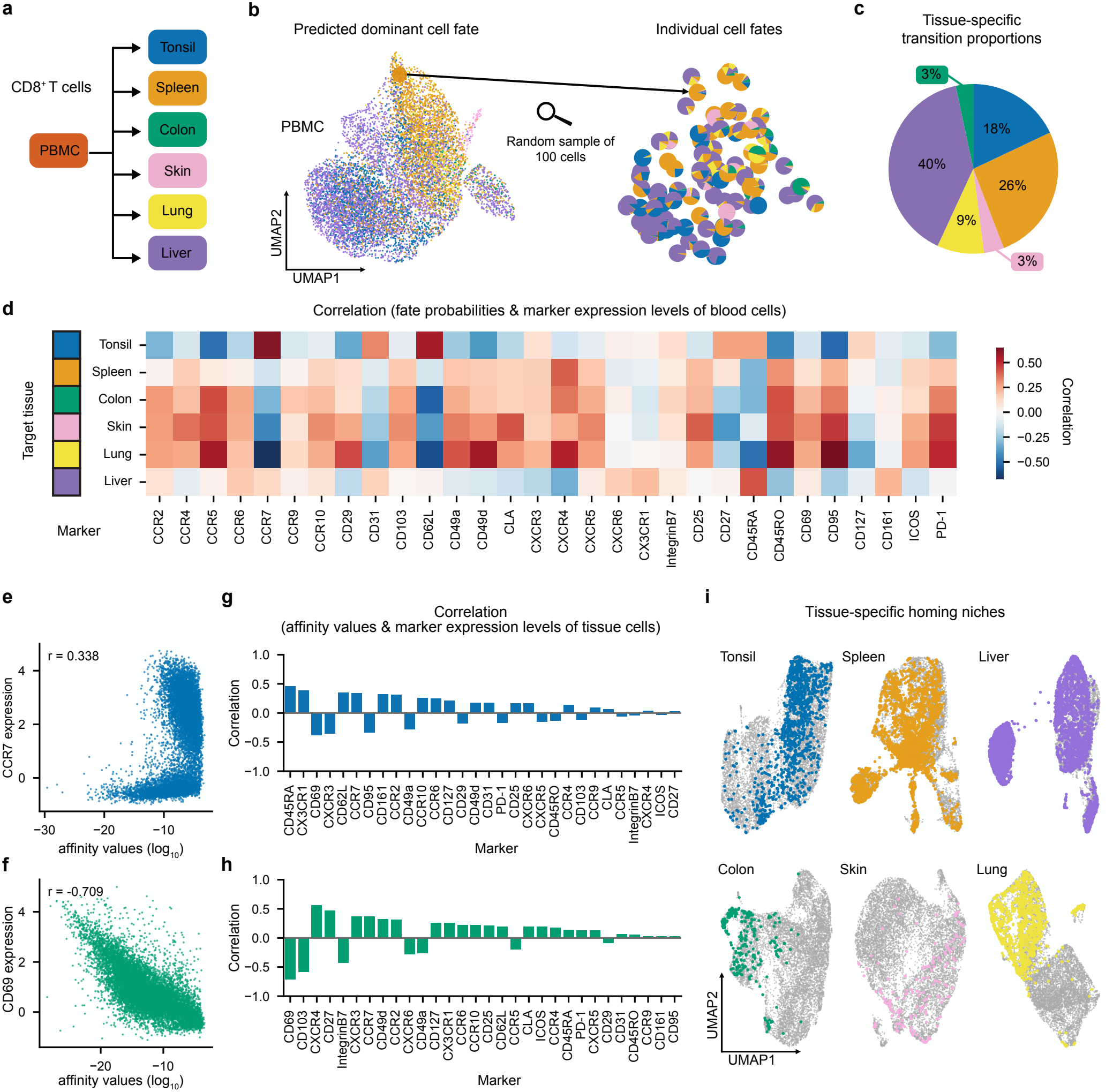
Cell fate prediction for CD8^+^ T cells based on mass cytometry data (Wong et al. 2016): (**a**) Tissue sites considered by Wong et al. [29] with PBMC measurements (=blood cells) used for fate prediction. (**b**) UMAP representation of 10^4^ CD8^+^ T cells of the PBMC coloured by the dominant fate prediction by *StitchOT* with individual fate predictions for 100 randomly sampled cells. (**c**) Predicted tissue-specific transition proportions for PBMC measurements. (**d**) Heatmap of the correlation of specific marker expression levels and predicted cell fates for the different tissues. (**e-f**) Exemplary correlation of marker expression levels for CCR7 and CD69 of individual tissue CD8^+^ T cells for the tonsil (**e**) and colon (**f**), respectively, with their affinity values towards blood cells. (**g-h**) Corresponding correlation values across the tissue-specific markers for tonsil (**g**) and colon (**h**). (**i**) Identification of homing niches of PBMC-CD8^+^ T cells with a high fate potential (*ψ* ≥ 1*/*3) to the particular tissue by projection into the UMAP representation of the CD8^+^ T cells measured at the respective tissue site. (**b-d**) are based on estimated cell fates by *StitchOT*, while analyses from the viewpoint of target cells (**e-i**) are based on *JointOT*. Correlations were computed using Spearman’s rank correlation.

Corresponding to previous observations, we identified tissue-specific trafficking signatures, with e.g. high expression of CCR7 and CD62L correlating with higher fate probabilities towards tonsil tissue, while being negatively correlated with entry into peripheral tissues (e.g. colon, skin) [32], as well as migration to the skin and colon being characterized by CCR4/CLA- and Integrin-*β7*/CD103-expression, respectively (Fig. 4d) [29, 33, 34]. Similar correlation patterns could be observed for CD4^+^ T cells for the particular marker/tissue combinations, with strong associations of CD62L and CCR7 expression for transition to the liver (Suppl. Fig. S4c).

Inferred affinity values for tissue cells, with high values characterizing cells that most likely recently entered the tissue, show distinct correlation behaviour with individual marker expression levels (Fig. 4e,f), characterizing tissue-specific homing patterns defined by combinations of beneficial and antagonistic marker expressions (Fig. 4g,h). For example, CD69-expression, a classical tissue-resident cell marker [35, 36], negatively correlated with the potential of having recently entered the tissue, which was consistently observed for CD8^+^ and CD4^+^ T cells across all sampled tissues (Fig. 4g,h, Suppl. Fig. S4e-h, Suppl. Fig. S5). In contrast, low CD103-expression of CD8^+^ T cells, but not of CD4^+^ T cells, in colon characterized their potential recent appearance corresponding to previous observations [37– 39], indicating tissue- and cell-type specific homing patterns. Further high and low marker expression levels indicating recent tissue appearance of CD8^+^ T cells involved classical trafficking receptors, such as CD62L/CXC3CR1 and CXCR6/CD49a, which are later down- and up-regulated, respectively, within tissue-resident cells [37].

Overall, this indicates that CellVADIS enables the determination whether blood cells would enter tissues within specific homing niches, i.e. areas within the marker space of the target tissue that correspond to recently entered cells. Projecting cells with a high fate potential into the corresponding tissue-specific marker spaces (see *Materials and Methods*) revealed that while peripheral tissues (i.e. lung, skin, colon) are characterized by specific homing niches, more heterogeneous patterns are observed for lymphoid tissues (tonsil, spleen) and the liver (Fig. 4i). In contrast, no such tissue-specific homing niches are predicted to exist for CD4^+^ T cells, except for tonsils, with more widespread projection patterns observed (Suppl. Fig. S4i). In summary, besides identifying trafficking signatures, CellVADIS enables the identification of ‘founder’ cell populations which could enable the determination of tissue-specific cell developmental pathways.

### Identifying tissue-specific trafficking signatures in transcriptomic data

With CellVADIS working for all different types of high-dimensional single cell measurements, we additionally applied our methods to data of the Cross-Tissue Immune Cell Atlas (TICA) containing scRNAseq-measurements of donor-matched tissues from 12 deceased donors [30]. Focusing on the T and innate lymphoid cell population as characterized in [30] comprising 20, 333 PBMCs, our method predicted that ~ 2% of the cells would migrate to gut-related tissues (i.e. jejunum), while transition to all other tissues (i.e. skin, spleen, lung, liver, bone marrow and lymph nodes) would be more or less evenly distributed (Fig. 5a,b). Restricting analysis to CD8^+^ T cells revealed comparable predicted transition proportions as obtained for the mass cytometry data (Fig. 4c,[29]), especially when accounting for potential unobserved tissue sites (Suppl. Fig. S6), while predictions for CD4^+^ T cells showed stronger deviations, independent of the approach used (Suppl. Fig. S6). Average tissue-specific cell fates for different cell types considered corresponded to the previously inferred relative abundances and significant enrichment of genes within specific tissues [30] (Fig. 5c).

**Figure 5:**
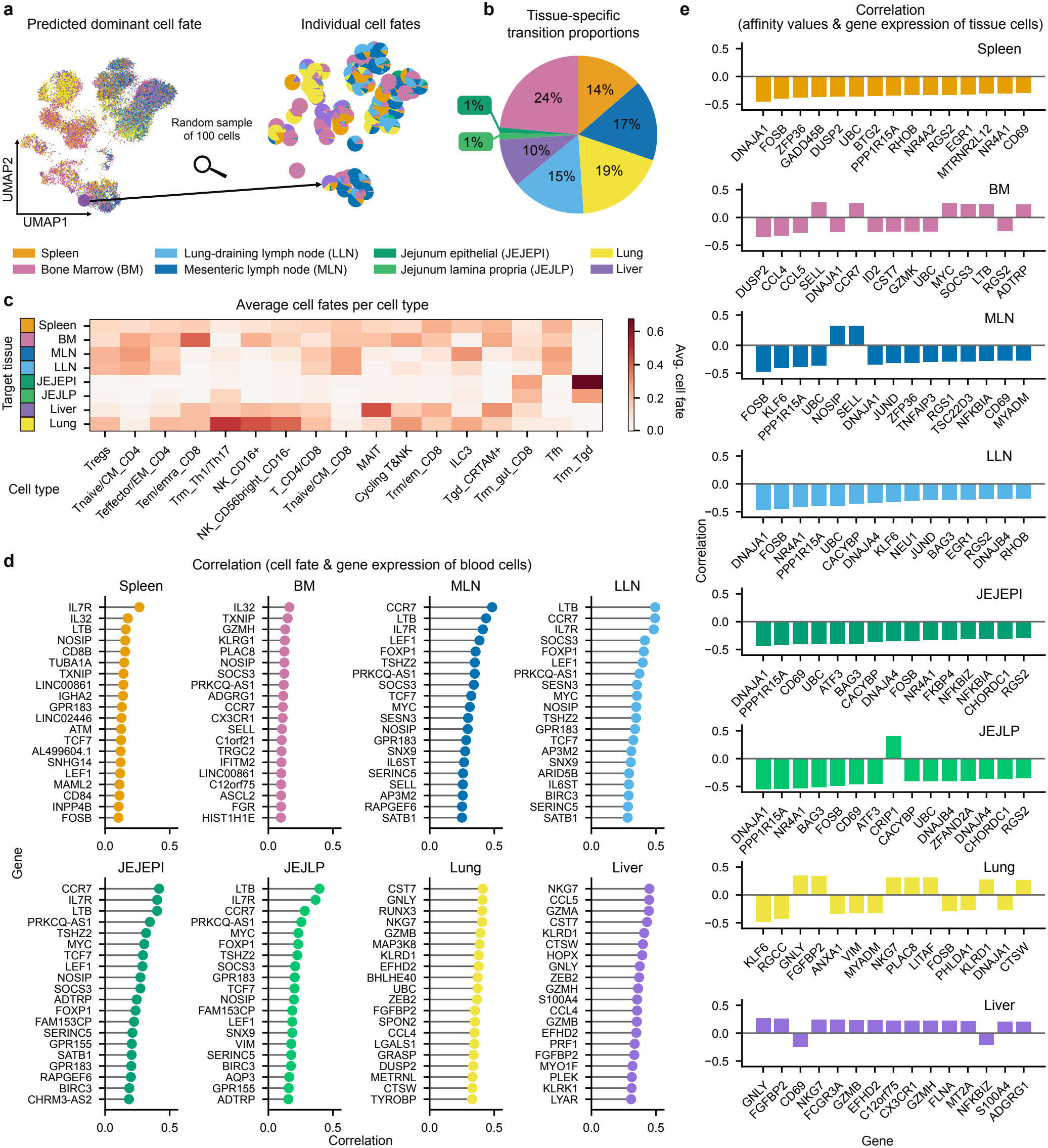
Cell fate prediction using scRNAseq-data (TICA, Conde et al. 2022): (**a**) UMAP representation of 20,333 T and innate lymphoid blood cells from the TICA-dataset coloured by the dominant fate prediction by *StitchOT* with individual fate predictions for 100 randomly sampled cells. (**b**) Predicted tissue-specific transition proportions for the cells in (**a**). (**c**) Average predicted cell fates of previously defined cell populations for the respective tissues. (**d-e**) Correlation of gene expression with predicted cell fate of blood cells (**d**) and affinity values for tissue cells (**e**) showing the top positively correlated 20 and the top 15 genes, respectively, for each tissue. (**e**) is based on the inferred OT-map by *JointOT*. Correlations were computed using Spearman’s rank correlation.

CellVADIS indicates tissue-specific gene expression patterns that correlate with cell transition potential, providing tissue-specific trafficking signatures based on the gene expression profile of individual blood cells (Fig. 5d). Known migration associated genes, such as CCR7 and SELL encoding CD62L, are observed to be positively correlated with confirmed migration fates, as e.g. to lymphoid tissues (e.g. BM, LN). Linking gene and corresponding marker expression generally indicated weaker correlations of tissue-specific transition probabilities with particular markers compared to relationships inferred from mass cytometry data (Suppl. Fig. S6). Despite the challenges in comparing the cross-tissue datasets by [29] and [30], marker-associated trafficking patterns for particular tissues seem to be robustly inferred for CD8^+^ T cells, while larger deviations between cell-fate prediction based on mass cytometry and scRNAseq-data are observed for CD4^+^ T cells (Suppl. Fig. S6).

Possible homing niches of blood cells (Suppl. Fig. S7) seem to be mostly characterized by antagonistic gene expression profiles (e.g. CD69), indicating their upregulation within tissues, with the inferred patterns potentially influenced by not restricting the analysis to trafficking associated genes, for which high expression levels would usually characterize recently entered cells (Fig. 5e). In general, tissue-specific trafficking signatures and identification of potential tissue entry points might differ dependent on the cell- and data-type used.

## Discussion

In this study, we presented CellVADIS as a computational framework based on optimal transport theory to infer cross-tissue cell trafficking dynamics. Being applicable to various high-dimensional single-cell measurements, our methods are able to predict cell fates and corresponding transition proportions of migrating cells accounting for unobserved tissue sites, as well as enable the identification of characteristic trafficking patterns for particular tissues.

In contrast to previous approaches that characterized tissue-specific trafficking patterns based on the expression level of known trafficking-associated markers or genes on cells within a particular tissue [29, 40, 41], our method infers cell signatures on cells that would most likely travel to the particular tissue from a given source tissue, i.e. the blood. Applying CellVADIS to mass cytometry and transcriptomic data from previously obtained cross-tissue atlases [29, 30] confirmed known tissue-marker associations. For example, CLA-expression was highly correlated with cell fate towards the skin, corresponding to previous observations [33, 34], with CCR2, CCR4, CCR5, CCR6, CCR9 and CCR10 expression additionally positively associated with migration towards peripheral tissues [42]. However, our analyses also indicated partly differing trafficking signatures and inferred transition proportions to particular tissues dependent on the cell and data type used. This could relate to the different types of measurements, i.e. measuring actual cell surface marker expression mediating cell entry vs. gene expression, as well as deviations in cell type and tissue matching for the considered datasets [29, 30]. In addition, variations in data pre-processing, i.e. considering the complete marker space vs. PCA-reduced gene expression patterns or choice of batch correction methods, could affect comparisons and identification of trafficking patterns. By normalizing expression levels according to relative marker distributions, one could neglect the necessity of absolute expression thresholds for specific markers to actually mediate cell entry into a particular tissue. In contrast, normalization of marker expression levels across cell populations [43–45], could affect the determination of individual relative cell fates. Further experimental approaches, as e.g. using cellular barcodes [46], could help to address this aspect.

Besides inferring characteristic trafficking signatures, CellVADIS enables the identification of homing niches within cellular marker spaces at which cells potentially enter a particular tissue. While CD8^+^ T cells were found to most likely enter peripheral tissues (such as lung, skin and colon) from the blood with a particular phenotypic profile, more heterogeneous patterns are observed in lymphoid tissues and the liver, corresponding to the higher blood perfusion in the latter. In contrast, CD4^+^ T cells were observed to express more heterogeneous homing patterns within the various tissues, corresponding to the observation that they express less CD69 and CD103 associated with tissue-resident phenotypes [35–37], which would impair the identification of particular homing niches within the phenotypic cell space. In general, CD69 is consistently inferred to be negatively correlated with the potential of cells to have recently entered the tissue, supporting the up-regulation of this marker to characterize tissue residency. By identifying potentially recently entered cells, CellVADIS supports the identification of tissue-specific cell differentiation pathways, with current trajectory inference methods requiring the definition of founder populations for identifying cell developmental pathways [21, 47, 48].

CellVADIS provides a novel approach to infer tissue-specific ‘postal codes’ based on cellular expression profiles, yet, might miss potential transition signatures. As inference of trafficking signatures relies on sufficient stable markers across tissues, rapid tissue-associated changes of entering cells, as e.g. the down- or up-regulation of particular markers inside tissues, could impair the identification of trafficking signatures and homing niches. This limitation cannot be addressed by either method without additional knowledge on the marker kinetics, with our model parameterized to match observed tissue-related dwell times of T cells [49]. Aiming to infer directed cell fates, variations in the methodological framework against synthetic data with known ground-truth also showed that CellVADIS is robust against the risk of actually mis-mapping ‘tissue-leaving’ cells ‘backwards’. In addition, general issues associated with data collection, including limited sampling, unequal cell distributions between tissues and unobserved tissue sites, could impair trafficking inference. However, although we assume 10^4^− 10^5^ cells per simulation to benchmark our method, sample sizes within the simulated tissues are significantly smaller and unevenly distributed (10^2^ −10^3^ cells), corresponding to actual experimental setups, showing that CellVADIS works robustly with additionally allowing to correct for unobserved tissue sites. Application to data with known biological ground truth of cell cross-tissue trafficking, as e.g. obtained by photoconversion [14] or cellular barcoding [46], could further help to validate CellVADIS and infer tissue-specific cell trafficking dynamics.

CellVADIS predicts cell trafficking fates from a pre-specified source compartment combining static snapshots of tissue samples from a common time point. While we used the blood as the source compartment in our analyses, applying CellVADIS with the assumption of different source compartments could help to identify cell circulation patterns and distinguishing tissue-specific circulating and resident populations. In general, *StitchOT* is preferred for cell fate predictions of source cells, while *JointOT* is better suited for analyses from the target cell perspective,e.g. to identify tissue-specific entry niches, due to their different ways of marginal relaxation. In comparison to other approaches looking at immune cell trafficking between different tissues [49, 50], our approach enables the inference of individual cell migration fate potentials rather than population-based migration rates by accounting for continuous marker expression profiles [20].

In summary, CellVADIS provides a unified framework for predicting immune-cell trafficking fates, identifying tissue-specific trafficking signatures and delineating candidate tissue-entry populations from cross-tissue single-cell snap-shots. Available as an open-source software package at https://github.com/GrawLab/CellVADIS, it supports the analysis of both protein expression and transcriptomic data. By linking circulating immune-cell profiles to potential tissue destinations, CellVADIS enables the generation of testable hypotheses about the establishment of tissue-specific immunity. These capabilities provide a foundation for identifying trafficking programs that could be targeted to direct immune responses towards specific tissues in vaccination and immunotherapy.

## Materials and methods

### Mathematical model to describe marker-driven cell trafficking dynamics

For method development and robust benchmarking based on synthetic data, we developed a stochastic, individual cell-based mathematical model in which individual cellular processes, such as cell differentiation and trafficking, are regulated and characterized by the continuous expression of cell-surface markers and chemokine receptors. Cells are localized in and traffic between different tissues, including the blood, lymphatic (e.g. spleen, lymph nodes, bone marrow) and peripheral tissues (e.g. gut) with the expression level of tissue-specific chemokine receptors determining the probability of cell migration into specific tissues. Individual marker expression levels are thereby described by stochastic up-regulation and degradation dynamics dependent on the actual cell location. The detailed description, implementation and parameterisation of the model, including model extensions, are given in Suppl. Information Text S1 and the accompanying tables, with the model code additionally provided at https://github.com/GrawLab/CellVADIS_analysis.

### Synthetic data generation

Simulations were performed by sampling parameters from uniform prior distributions by mainly varying parameters that are assumed to alter migratory behaviour (see Tab. S1, S2, S3, S4). For each simulation, a total of 10^4^ −10^5^ cells were simulated and followed over 30h in simulated time after considering a burn-in period of 48h to allow for system calibration. At 24h, the blood and various tissue sites were sampled, and the different OT methods were applied to the blood cell data. Hereby, only simulated blood cells that were observed to actually transit to another tissue within 6 hours after sampling were considered in the analysis. This usually comprises all cells as ~ 99% of simulated cells leave the blood within minutes after entering, corresponding to biological observations [51]. Simulations are further filtered to only contain robust simulations with sufficient cell sizes where every location contains at least 500 cells, and at least 100 blood cells have a true next location corresponding to each target tissue. Performing a total of 10^4^ simulations, after filtering, 4233 simulations were considered for further analysis.

To determine method robustness in case of an unobserved tissue *U*, sensitivity was analysed using combinations of the steepness parameter *γ* (Eqs. (44)-(45)) and the fraction of cells expressing marker ⋀ (*λ* ^⋀^ (see Eq. (25) and Suppl. Table S3), considering 10 parameters each within a 2D parameter grid, given log-spacing in [10^*−*3^, 1] (*γ*) and linear-spacing in [0, 1] (*γ* ^⋀^), respectively. For each pair of (*γ, λ* ^⋀^), the model was simulated utilizing the same 1000 parameter combinations drawn from the uniform prior distributions as before. Simulations and considered blood cells were filtered as indicated above.

### Optimal transport for inferring cell trafficking

To determine cell trafficking dynamics between tissues we developed novel methods based on optimal transport. Consider two cell populations of size N and M, with each cell characterized by the intensities of D different markers. The corresponding marker profiles of the two cell populations are represented by matrices *X* ∈ℝ^*N ×D*^and *Y* ∈ℝ^*N ×D*^, respectively. Each cell is additionally assigned a weight, so called marginals, *a*_*i*_ and *b*_*i*_, such that 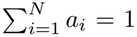 and 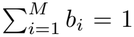 These weights can be chosen to represent cellular turnover [27, 28], i.e. cells with weight *a*_*i*_ < 1/*N* (*b*_*i*_ < 1/*M*) indicate cells that are likely to disappear while cells with *a*_*i*_ > 1/*N* (*b*_*i*_ > 1/M) potentially proliferate. If prior knowledge of these quantities is lacking, all cells are assigned uniform weights, i.e. *a*_*i*_ = 1/*N* for *i* = 1, …, *N* and *b*_*i*_ = 1/*M* for *i* = 1, …, *M*. Optimal transport aims to find a mapping *T* ∈ℝ^*N ×M*^ that links the cellular profiles between the two groups with *T*_*ij*_ representing the amount of mass sent from source cell i to target cell j. The OT-map T additionally needs to satisfy the marginal constraints *T* ×**1**_*M*_ = *a* and T ^*⊤*^ *×***1**_*N*_ = *b*, with **1**_*M*_ = (1, …, 1) ^*⊤*^ ∈ℝ^*M*^, as well as minimize a cost matrix *C* ∈ℝ^*N ×M*^, with the entries *C*_*ij*_ defined by a suitable cost function that relates marker profiles of source to target cells. For simplicity the euclidean distance, i.e. *C*_*ij*_ = *c*(*x*_*i*_, *y*_*j*_) = ||x_*i*_ *−* y_*j*_||_2_, with *x*_*i*_, *y*_*j*_ ∈ℝ^*D*^, is often used, although the choice could depend on the data modality. Thus, the objective function reads

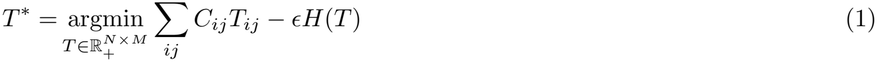

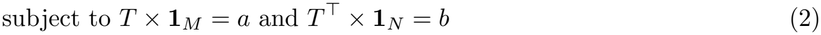

with *H*(*T*) = −*Σ* _*ij*_ *T*_*ij*_ log(*T*_*ij*_) defining the entropy regularizer term, which ensures that the objective function is strictly convex and allows the use of efficient optimization algorithms [52]. The parameter *ϵ*controls the degree of entropic regularization, with larger values yielding a more diffuse OT-map T and greater mass splitting across target cells, while smaller values favor more concentrated transport assignments [28, 52]. Eq. (1) relates to balanced optimal transport, for which the solution *T*^*^ is enforced to exactly match the predefined marginals a and b. In scenarios where exact enforcement of the marginal constraints is too restrictive, small variations are allowed by using unbalanced OT [31] to account for mass fluctuation between the two distributions by mass gain and loss, as e.g. due to cell proliferation, biased sampling, or cell differentiation within tissues. Here, we replace the exact constraints *T* ×1_*M*_ = a and *T* ^⊤^ × 1_*N*_ = *b* by soft Kullback-Leibler constraints, which change Eq. (1) to

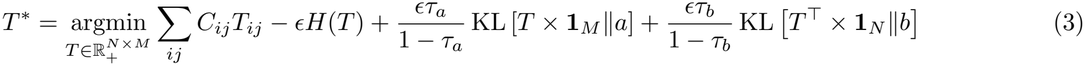

The parameters *τ*_*a*_, *τ*_*b*_ *∈* (0, 1) are thereby weighting parameters related to the marginals a and b, respectively (see also [28]), with *τ*_*a*_, *τ*_*b*_ *⟶* 1 recovering the exact marginal enforcement.

To infer the potential of cells within the blood to migrate to either of the specific L other tissues (e.g., LN, gut, and spleen), we therefore need to solve an optimal transport problem that simultaneously maps the cell distribution within the blood to the cell distributions within the individual target tissues. Thereby, we developed two different approaches, which either treat this mapping as separate or combined OT-problems.

### StitchOT - combining independent OT-maps

Assuming that a cell from the blood can migrate to L different possible locations, the mapping of cells between these different locations can be treated as L separate unbalanced OT problems. *StitchOT*, therefore defines a cost matrix 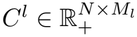 based on the L2-norm between the blood and non-blood cells in phenotypic marker space, with M_*l*_ defining the cell population size in location *l* ∈ {1, …, *L*}. Without further knowledge, the marginals 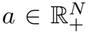and 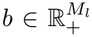 are then considered to be uniformly distributed with

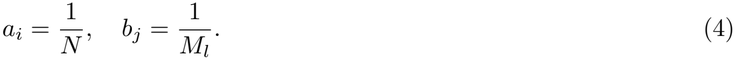

For each location *l* ∈ {1, …, *L*}, an OT-problem is then solved separately to determine the OT-map 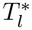 corresponding to Eq. (3):

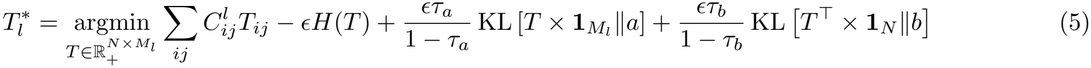

Hereby, we relax the blood marginal constraint by setting *τ*_*a*_ = 0.5, such that the marginal weights of the blood cells can be interpreted as an affinity rate of each blood cell to migrate to the specific location *l*. In contrast, the marginals for the target tissues are enforced more strictly, as most cells within the tissue are assumed to have infiltrated from the blood circulation at some point. However, to account for localized phenotypic changes, as well as tissue retention, the marginal constraints for these tissues are also slightly relaxed, setting *τ*_*b*_ = 0.995.

Since each transport problem is solved separately, the resulting transport matrices do not directly encode the relative contribution of the different target tissues. We therefore introduce a tissue-specific weighting factor *ω*_*l*_ before combining the individual transport matrices by considering the fraction of blood cells that increase their marginal value above the initial uniform reference marginal of 1/*N*, with

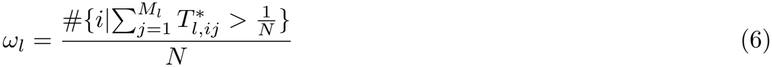

These weights are normalized such that Σ_*l*_ω_*l*_ = 1. The global OT-map 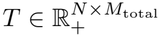, with 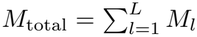 is then obtained by “stitching” together the weighted location-specific OT-maps 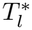, i.e.

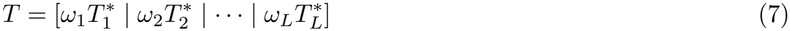

The weighted transport assignments are subsequently aggregated by target tissue and normalized across tissues for each blood cell, yielding the fate matrix *F ∈* ℝ^*N L*^, with

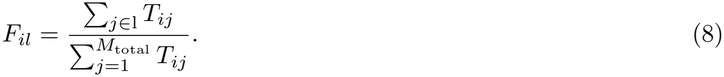

Each entry *F*_*il*_ represents the inferred relative trafficking fate of blood cell *i* towards tissue *l*, with 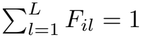.

#### JointOT - simultaneous optimization of global OT-map

In contrast to *StitchOT, JointOT* integrates all *L* target tissues into a single OT problem and determines a joint OT-map between the blood and all target cells. If 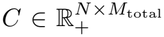 defines the cost matrix, then the optimization problem relates to finding the joint OT matrix *T* according to

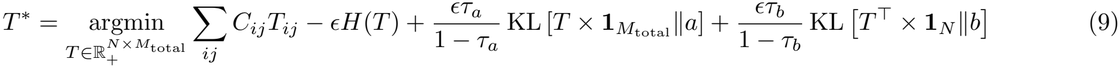

analogous to Eq. (5). In contrast to *StitchOT*, for *JointOT* we relax the marginal constraint on the non-blood cells, i.e. setting τ_*b*_ = 0.5 while enforcing the uniform mass constraint on the blood cells more strictly (τ_*a*_ = 0.995). Thus, each blood cell contributes approximately its prescribed mass, while the amount of transport mass assigned to individual non-blood cells can vary to minimize the overall transport cost. By aggregating the transport mass assigned to target cells according to their tissue identity, we directly obtain the fate matrix 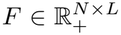 as defined in Eq. (8). Both methods rely on the Python Optimal Transport (POT) package [53, 54] for computing the raw optimal transport outputs.

### OT-method extension to account for unobserved tissues

In general, optimal transport will characterize mis-mapped cells by a high cost associated with a mapping to any of the sampled tissues. Therefore, we adapted our methods to account for the probability of a cell to be mis-mapped.

### StitchOT

For *StitchOT* we extended the fate matrix 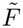 by the probability of a cell to be mismatched, *F*_*i*(*L*+1)_, i.e. to not migrate to any of the sampled tissue sites *l* = 1, …, *L*, based on the inferred marginal values 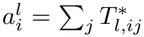 with

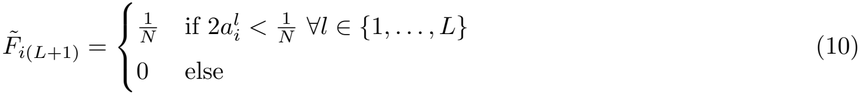

Hereby, *N* denotes the total number of blood cells. Thus, if the predicted probabilities for a cell to be mapped to any sampled tissue will be too small, the dominant cell fate after normalizing the fate matrix 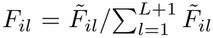 will indicate a mis-mapped cell belonging to the unobserved tissue.

#### JointOT

For *JointOT*, the adaptation is similar to *StitchOT*, with the assigned unobserved probability determined based on the mass deficit of each blood cell compared to the reference marginal 1*/N*:

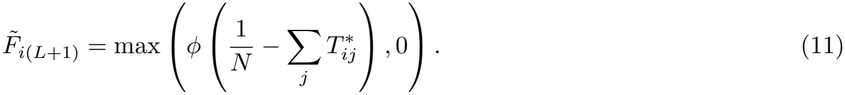

This way, blood cells with no marginal deficit or an increased inferred marginal will receive an unobserved probability of 0, while cells that have a positive marginal deficit increase their unobserved fate potential. As *JointOT* strongly penalizes mass destruction for source cells (due to *τ*_*a*_ = 0.995), the mass deficit will never reach its theoretical maximum of 1/*N*. Therefore, we multiply the raw mass deficit by a hyperparameter *ϕ* to increase the practical mass deficit. Hereby, we choose *ϕ* = 100. By adding the additional dimension to the fate matrix 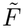 before normalizing, the assigned probability is independent on the scale of the marginals (i.e. number of blood cells), as 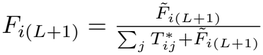

To quantify the robustness of the methods given unobserved tissue sites, the different metrics introduced below were applied to the full fate matrix, i.e. assuming that the unobserved tissue site *U* could be sampled, as well as the conditional fate matrices only considering blood cells not trafficking to *U*. In the latter case, the metrics are computed on the re-normalized and reduced fate matrix *F*_*il*_ = *F*_*il*_/(1 *-F*_*i*(*L*+1)_) with *l* ∈ {1, …, *L*} and 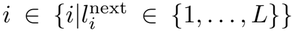 whereby 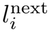 defines the true next location of cell *i*. If *U* is not observed and no correction is applied, the full fate matrix *F* is defined with *F*_*i*_(*L* + 1) = 0.

### Assessing method performance

#### Accuracy

The total accuracy, ACC, is defined as the fraction of correctly predicted locations across all blood cells, *N*, with the predicted location for each cell *i* defined as the location with the highest predicted migration probability based on the fate matrix *F* derived from the OT-map:

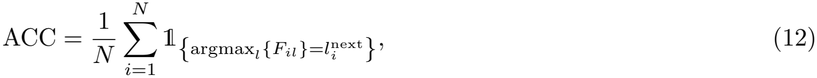

with 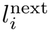 defining the true next location of cell *i*. Given that the actual cell fate within the simulated ground truth data is sampled from a multinomial distribution determined by the marker-driven migration probabilities associated with each tissue, the total accuracy defined by Eq. (12) represents a rather conservative metric, i.e. ignoring the stochasticity influenced by the actual values and relative differences between *F*_*il*_,*l ∈ {*1,…, L*}*. Therefore, we also defined a *threshold accuracy, ACC*_*ψ*_, by assuming that the fate is correctly predicted if the transition probability to the true next location 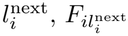 is larger than a specified threshold *ψ ∈* (0, 1), i.e.

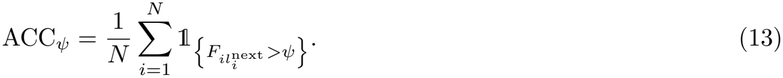

The *average threshold accuracy*, 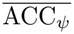, was then determined by integrating *ACC* across all possible thresholds *∈ ψ* (0, 1), i.e.

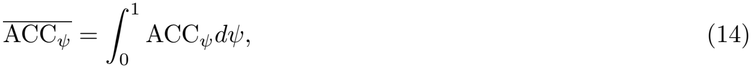

to account for sensitivity due to arbitrarily chosen thresholds.

#### Relative accuracy

As the true location fate for each simulated cell is determined by randomly sampling from a multinomial distribution, the accuracy for predicting the location fate of an individual cell is not expected to reach 1, even when knowing the exact tissue-specific migration probabilities of each cell. Therefore, the different OT methods will not perform better than a *practical optimal accuracy* achieved by considering the true migration rates calculated for each cell and tissue (see Eq. (16)). As the standard measures of accuracy (see above) would therefore underestimate the true accuracy of the different methods, we define the *relative average threshold accuracy*, 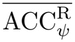, as the main metric for assessing method performance. 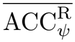 normalizes the average threshold accuracy 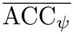, which relies on the predicted transition values *F*_*il*_ of the OT-methods, by the average threshold accuracy defined by the ground truth values 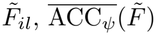, of the simulation, thus,

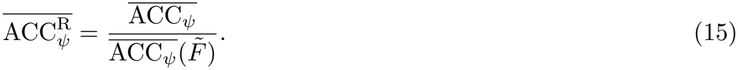

Here, the ground truth fate matrix 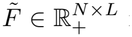 is determined by calculating the cell-specific migration rates based on the individual marker expression levels for all blood cells (see Eqs. (33)-(36) in Suppl. Text S1), with

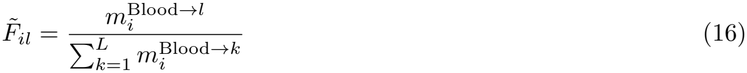

### Integrated absolute error

For specific thresholds ψ, the accuracy *ACC* _*ψ*_ of the method could be higher than the ground truth threshold accuracy, 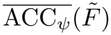. In this case, the mapping would be overconfident, resulting in 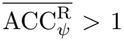 for individual *ψ ∈* (0, 1). In correspondence to Eq. (14), we therefore also determined the integrated absolute error of the threshold accuracy, defined by:

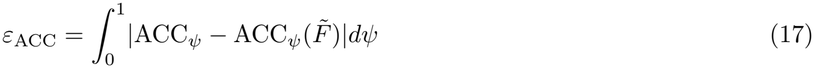

as an additional measurement to assess method performance accounting for overconfidence.

#### Fate matrix comparison

To further assess method performance directly based on the fate matrix *F*, we determined the extent to which the inferred cell-based fate probabilities deviate on average from the ground truth transition probabilities for each location *l* ∈ {1,…, *L*} across all blood cells. This *mean total variation distance*, Δ_F_, is then defined by:

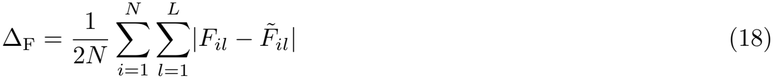

#### Tissue-specific transition proportions

With OT-methods predicting cell trafficking from blood to specific tissues, an additional quantity to assess overall method performance relates to the ability to correctly predict the proportion of blood cells that would end up in a specific tissue site, i.e. defined by the *tissue-specific transition proportions*. For the ground truth simulation, the fraction of blood cells trafficking to tissue *l* = 1,…, *L* is given by 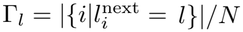, with N denoting the total number of blood cells, and *Σ* _*l*_ *Γ*_*l*_ = 1. The corresponding predicted transition proportions are then computed based on the normalized fate matrices of the inferred OT-maps according to

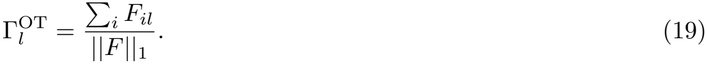

Note here that ||*F*||_1_= *N*, if the full fate matrix is considered. Method performance is then validated by comparing the deviations of *Σ*= *{* _*l*_*}*_*l*=1,…,*L*_ and 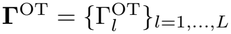 with Δ_Γ_ = |Γ − Γ^OT^|/2.

## Data preparation - Wong et al. Immunity 2016

FCS-files for pre-processed and defined CD4^+^ and CD8^+^ T cells were obtained from FlowRepository [55], under Repository-ID FR-FCM-ZZTM. All pre-transformed FCS-files from the 30 independent tissue samples, including adult blood, tonsil, spleen, colon, skin, lung and liver (see [29] for details), were pooled across tissues and donors using Pytometry [56]. To account for potential batch effects between donors and tissues, we computed gene-wise z-scores, which shifts the representation from absolute expression levels to relative marker profiles. Biologically, this removes scaling artifacts as introduced through staining intensity variations between tissues or instrument sensitivity. After normalization we sub-sampled 10^4^ cells of each tissue for further analysis. PBMC cells from the adult blood were selected as source (blood) cells with the remaining tissues treated as target tissues. Markers that were classified as differentiating, activating, or trafficking according to the categories used in the original publication were considered within the analyses.

## Data preparation - Conde et al. Science 2022

All analyses were conducted on the pre-processed dataset for T and innate lymphoid cells from the original study in [30] and obtained from *https://www.tissueimmunecellatlas.org*. Analyses were performed in Python, using `Scanpy` [57] and `AnnData` [58] for further data processing, before conducting analysis within CellVADIS. After loading the data, we calculated highly variable genes (`sc.pp.highly variable genes`) using the donor information as batch key. A 50-dimensional PCA data reduction (computed using `sc.tl.pca`) was used as input for batch integration using `Harmony` [59] in `Scanpy` (`sc.external.pp.harmony integrate`). The resulting 50-dimensional batch-corrected PCA coordinates were used for our OT-based analysis using CellVADIS.

Only tissue samples that contained at least 1000 cells were considered within the analysis (i.e. including PBMC, spleen, bone marrow (BM), lung-draining lymph nodes (LLN), mesenteric lymph nodes (MLN), jejunum epithelium (JEJEPI), jejunum lamina propria (JEJLP), lung, and liver), with subsequently subsampling 5000 cells for all target tissues, i.e. all tissues except PBMCs. PBMCs considered as the source (blood) cells were not sub-sampled in order to increase robustness in estimating the average cell fate for rare cell types, leaving a total of 20,333 source cells for cell fate prediction.

After computing cell fates using CellVADIS, downstream analyses were performed on the highly variable genes, excluding Ribosomal (RPS, RPL), mitochondrial (MT-), heat shock (HSP), and HLA genes to prevent them from dominating gene-wise analyses.

For cell-type specific analysis we used the manually curated cell type definition as provided by Conde et al. [30]. To compare to [29], cell populations were combined according to CD4=*{*Treg, Teffector/EM CD4,Tnaive/CM CD4,Tfh, Trm Th1/Th17*}* and CD8=*{*Tem/emra CD8, Tnaive/CM CD8,Trm/em CD8, Trm gut CD8*}* with different tissue sites compared as shown in Suppl. Fig. S6.

## Identification and visualisation of homing niches

To determine if recently infiltrated cells enter tissues in specific homing niches defined by characteristic marker combinations, we consider the target marginals b_*i*_ = *T*^⊤^ × 1_*N*_ as estimated by *JointOT* for all cells inside target tissue *l*, which are denoted by *ℐ*_*l*_ = *{i*|*l*_*i*_ = *l*}. The reduced tissue-specific marginal vector 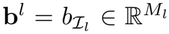 is then correlated with the corresponding expression vector 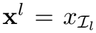 for the specific marker or gene x using Spearman’s rank correlation.

To help visualize these potential tissue entry points, we projected blood cells that were predicted to migrate to the particular tissue based on the OT-methods into the UMAP representation of the sampled tissue population using a weighted average of the tissue coordinates. To this end, let 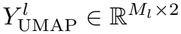 define the UMAP-coordinates of all *M*_*l*_ cells in tissue *l*. Only focusing on blood cells, for which the predicted fate probability for the particular tissue is larger than a threshold *θ*, i.e. S_*l*_ = *{*(*i, j*)|*F*_*il*_ > θ and j ∈ *ℐ*_*l*_*}*, we determine a coupling matrix 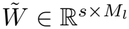 as a subset of the OT-map *T*, with *w*_*ij*_ = *t*_*ij*_ for all (i, j) ∈ *S*_*l*_ and s = |*{*i|(i, j) ∈ S_*l*_*}*|. Normalizing 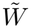 across the rows, we obtain the weighting matrix W with

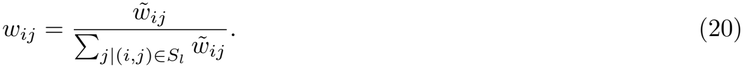

The projected marker coordinates of the blood cells with *F*_*il*_ > θ, and θ = 1/3, within the tissue-specific UMAP-projection are then defined by

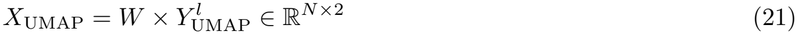

with N defining the total number of blood cells. If these cells cluster within the tissue-specific marker space, this would suggest the existence of homing niches that characterise recently entered cells within the particular tissue. For this analysis, *JointOT* is preferred as relaxing the marginal constraints produces larger mass variations among non-blood cells, yielding a more heterogeneous weighting matrix W.

## Acknowledgments

We thank Julia Hitschfel, Pascal Lukas, Simon Völkl, Anna-Lena Rösch and Wolfgang Kastenmüller for critical reading of the manuscript. This work was supported by the Boehringer Ingelheim Foundation within the RiseUp!-program (project RETENTION) and the Hightech Agenda Bavaria to F.G.. The authors gratefully acknowledge the scientific support and HPC resources provided by the Erlangen National High Performance Computing Center (NHR@FAU) of the Friedrich-Alexander-Universität Erlangen-Nürnberg (FAU). The hardware is partially funded by the German Research Foundation (DFG). The funders had no role in study design, data collection and analysis, decision to publish or preparation of the manuscript.

## Author contributions

Conceptualization: F.G.; Method development: L.W., S.V., F.G.; Software implementation: L.W.; Development of mathematical model for benchmarking: L.W., S.V., F.G.; Data curation: L.W., Formal analysis: L.W., F.G.; Investigation: L.W., S.V., F.G.; Data interpretation: L.W., S.V., J.H., F.G.; Visualization: L.W., F.G.; Writing - original draft: L.W., F.G.; Writing - review: all authors.

