## Supplementary Information for "CellVADIS: Inferring cross-tissue immune cell trafficking dynamics from single-cell snapshots"

**Text S1:** An individual cell-based model to describe marker-driven cell trafficking

**Text S2:** Extended explanations for unobserved tissue sites and cell fate prediction

**Text S3:** OT-method variations to assess method robustness

**Figure S1:** Mathematical model of marker-driven cell trafficking dynamics

**Figure S2:** Accuracy of cell fate prediction according to variations in the methodological framework

**Figure S3:** Extended metrics for method assessment given unobserved tissues

**Figure S4:** Cell fate prediction based on mass cytometry data for $CD4^+$ cells (Wong et al. 2016)

**Figure S5:** Trafficking signatures for different tissues and cell types (Wong et al. 2016)

**Figure S6:** Comparing cell fate prediction based on different data types

**Figure S7:** Prediction of homing niches for T & innate lymphocytes based on the cross-tissue immune atlas (TICA)

**Table S1:** General simulation parameters

**Table S2:** Marker-specific expression and degradation rates

**Table S3:** Parameters accounting for cell-to-cell variability in marker expression

**Table S4:** Parameters describing tissue-specific migration dynamics

#### Text S1: An individual cell-based model to describe marker-driven cell trafficking dynamics

For method development and robust *in silico* testing, we developed a stochastic, individual cell-based mathematical model. Cellular processes, such as cell differentiation and trafficking, are regulated and characterized by the continuous expression of cell-surface markers and chemokine receptors on individual cells, with cells localized in and trafficking between different tissues, including the blood, lymphatic (e.g. spleen, lymph nodes, bone marrow) and peripheral tissues (e.g. gut). Thereby, the expression level of tissue-specific chemokine receptors determines the probability of cells to migrate into specific tissues (Suppl. Fig. S1a). Individual marker expression levels follow stochastic up-regulation and degradation dynamics dependent on the actual cell location (Suppl. Fig. S1b) influencing cellular distributions (Suppl. Fig. S1c) and enabling the generation of individual cell-based measurements as obtained experimentally (Suppl. Fig. S1d). The individual processes of cell differentiation, stochastic marker regulation and cell trafficking dynamics of the multi-scale mathematical model are thereby described as follows:

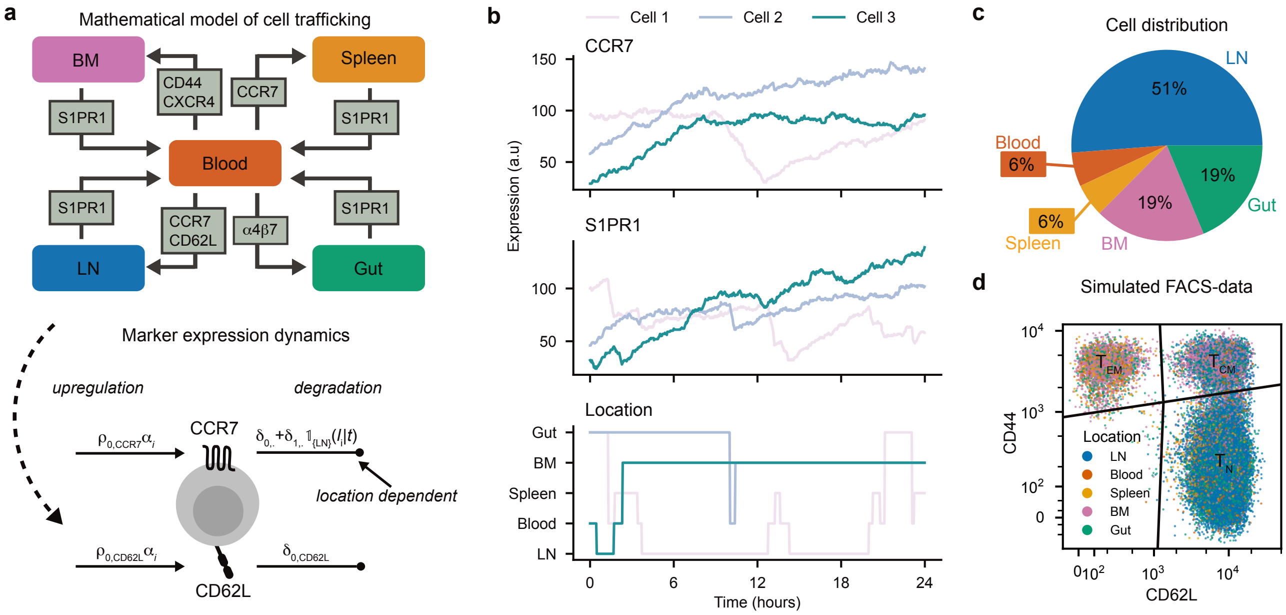


**Figure S1: Mathematical model of marker-driven cell trafficking dynamics:** (**a**) Sketch of the individual cell-based mathematical model developed to simulate cell trafficking dynamics considering the blood as the central transit compartment. Cellular marker expression profiles determine cell infiltration and egress dynamics with particular locations influencing marker upregulation and degradation. (**b**) Simulated dynamics of CCR7- and S1PR1-expression of three individual cells in correspondence to their locations over time. (**c**) Cell distribution between different organs at a particular time point with (**d**) exemplary FACS-plot showing the ability to mimic identification of particular T cell phenotypes based on CD44 and CD62L marker expression as obtained within experimental data. Colors indicate the location of each cell at the time point of sampling.

**Cell differentiation & marker expression dynamics:** Continuous marker expression dynamics of a cell $i$ are determined by a set of reactions, given in the following general form:

$$\frac{dX_i}{dt} = k_{i,x} - \gamma_{i,x}(l_i|t)X_i(t) \tag{22}$$

with $X_i(t)$ representing the expression level of marker $x$ on cell $i$ at time $t$, and $k_{i,x}$ and $\gamma_{i,x}$ defining the cell-specific expression and degradation rates, respectively, dependent on the cell's current location $l_i$ at time $t$. Neglecting the index $x$ in the following to simplify notation, in the most general form, $k_i$ and $\gamma_i$ are defined as a linear combination of the baseline rates, i.e. $\rho_0$ and $\delta_0$, and the location-specific contributions according to

$$k_i = \rho_0 \alpha_i \tag{23}$$

$$\gamma_i (l|t) = \delta_0 + \delta_1 \mathbb{1}_{\mathcal{L}}(l_i|t), \tag{24}$$

with $l_i$ defining the tissue cell $i$ is currently in and the indicator function $\mathbb{1}_{\mathcal{L}}$ being 1 if $l_i$ relates to a specific tissue in the set of locations $\mathcal{L} \subset \{1, \ldots, L\}$ which influence the expression of the particular marker and 0 otherwise. In order to account for cell-to-cell variability and intrinsic noise in flow cytometry measurements, the baseline expression rate $\rho_0$ is additionally multiplied by marker-specific random factors $\alpha_i$ that are drawn from a probability distribution defined by

$$\alpha_i \sim \lambda \text{Lognormal}(\mu_L, \sigma_L) + (1 - \lambda)\mathcal{N}(\mu_G, \sigma_G), \tag{25}$$

with $\mu_G > \mu_L$, such that the resulting distribution captures the frequently observed bimodality of marker expression levels in flow cytometry data. Thereby, the parameters $\lambda$, $\mu_G$, $\sigma_G$, $\mu_L$ and $\sigma_L$ are marker specific and can be adjusted to control the degree of bimodality and variability in the marker expression across cells, with $\mu_G$ and $\sigma_G$ chosen to mitigate potential values of $\alpha_i < 0$, which are otherwise set to 0 (see Suppl. Fig. S1d for an example of the bimodal marker distribution of the markers of CD44 and CD62L). Hereby, $\gamma_i$ does not include a random cell-specific parameter $\alpha_i$, as cell heterogeneity is covered by $k_i$.

Our model currently accounts for markers addressing cell differentiation status (CD44, CD69) and tissue-specific homing dynamics (CCR7, CD62L, S1PR1, CXCR4, $\alpha 4\beta 7$). Thereby, we distinguish between markers for which the expression dynamics is influenced by the cell location, and those that are regulated independently of it. A detailed description of the marker-specific reaction kinetics is given in the following, with their individual parameterizations given in Tables S1-S4.

**(a) Markers not being actively affected by cell location:** We currently assume that several markers are not actively regulated by cell trafficking and its change of location, or cell activation and proliferation status. In this case, Eq. (23) simplifies to $k_i = \rho_0 \alpha_i$ and $\gamma_i = \delta_0$. This includes the following markers:

- **CD44** is a cell activation marker that is permanently upregulated after T-cell activation and often used to distinguish naive (CD44 low) from activated/memory (CD44 high) T cells. Currently, CD44 expression is kept at the cell-specific steady state level determined by the random factor $\alpha_i$.
- $\boldsymbol{\alpha 4\beta 7}$ is an integrin that promotes homing to the gut by mediating adhesion to the gut endothelium [1, 2]
- **CD69** is an early activation marker that is often used to identify tissue resident T cell populations, as it inhibits S1PR1 expression on the cell surface and thereby promotes tissue retention [3, 4].
- **CD62L**, or L-selectin, is essential for T cell rolling on high endothelial venules, which are present within lymph node paracortical regions, promoting homing to lymphatic tissue [5].

Individual cell marker expression levels will still randomly fluctuate due to the stochastic simulation of marker regulation.

**(b) Markers actively affected by cell location:** For other markers that actively change their expression levels dependent on the cellular location, Eq. (23) is defined as follows:

**CCR7** is a chemokine receptor that is expressed on naive and central memory T cells and promotes homing to lymph nodes by mediating chemotaxis towards the CCR7 ligands CCL19 and CCL21 [5, 6], which are highly abundant in secondary lymphoid organs. In our model, the expression of CCR7 is downregulated in secondary lymphoid organs (lymph nodes and spleen, i.e. $l \in \{\text{LN,Spleen}\}$) in a time-dependent manner, which mimics the receptor internalization upon ligand binding. After exit from the lymphoid organs, CCR7 expression is restored. Thus, the individual rates are defined as follows

$$k_i = \rho_0 \alpha_i \tag{26}$$

$$\gamma_i(l_i|t) = \left(\delta_0 + \delta_1 \mathbb{1}_{\{\text{LN,Spleen}\}}(l_i|t) \frac{\Delta_i^\nu}{T_{\text{CCR7}}^\nu + \Delta_i^\nu}\right) \tag{27}$$

where $\mathbb{1}_{\{\text{LN,Spleen}\}}$ is 1, if cell $i$ is in either the lymph node or the spleen at time $t$. $\Delta_i$ denotes the time of cell $i$ since tissue entry.

**CXCR4** is a chemokine receptor that promotes homing to the bone marrow (BM) by mediating chemotaxis towards its ligand CXCL12, which is highly abundant in the BM. Like other G-coupled protein receptors, CXCR4 is internalized upon ligand binding [7, 8], which is integrated in the model by increasing the degradation rate of CXCR4 in BM. Upon egress, CXCR4 expression is restored, leading to the following equations:

$$k_i = \rho_0 \alpha_i \tag{28}$$

$$\gamma_i(l_i|t) = \left(\delta_0 + \delta_1 \mathbb{1}_{\{\text{BM}\}}(l_i|t) \frac{\Delta_i^\nu}{T_{\text{CXCR4}}^\nu + \Delta_i^\nu}\right) \tag{29}$$

**S1PR1** binds to its ligand S1P, which is highly abundant in the blood and promotes egress from lymphoid organs. Resident memory T cells in peripheral tissues express CD69 which inhibits S1PR1 expression and thereby promotes tissue retention [9]. In the blood, S1PR1 is internalized to promote fast reentering to tissues [6, 10–12].

$$k_i = \rho_0 \alpha_i \tag{30}$$

$$\gamma_i(l_i|t) = \left[\delta_0 + \delta_1 \left(\mathbb{1}_{\{\text{Blood}\}}(l_i|t) \frac{\Delta_i^\nu}{T_{\text{S1PR1}}^\nu + \Delta_i^\nu} + \frac{\text{CD69}_i^{\nu_{\text{CD69}}}(t)}{K_{\text{CD69}}^{\nu_{\text{CD69}}} + \text{CD69}_i^{\nu_{\text{CD69}}}(t)}\right)\right] \tag{31}$$

**Cell trafficking dynamics:** Cell migration to tissues is regulated by the expression of specific chemokine receptors (CCR7, CXCR4, S1PR1) and their interaction with their corresponding ligands (CCL19, CXCL12, S1P), as well as by the expression of adhesion molecules (CD44, CD62L, $\alpha4\beta7$) (Suppl. Fig. S1a). In our model, the trafficking of cells between different tissues $A$ and $B$ is probabilistically determined by the individual expression levels of these markers on each cell. Distinguishing between the expression levels of markers that would promote the trafficking of the cell from tissue A to B, $X_{i,m}$, $m \in \mathcal{M}_{\{A,B\}}$, and those that prevent the cell from leaving, $X_{i,d}$, $d \in \mathcal{D}_A$, we define the migration rate of cell $i$ from tissue $A$ to tissue $B$ by:

$$m_i^{A\to B} = \lambda_{A\to B} \prod_{m \in \mathcal{M}_{\{A,B\}}} \frac{X_{i,m}^\nu(t)}{K_m^\nu + X_{m,i}^\nu(t)} \prod_{d \in \mathcal{D}_A} \left(1 - \frac{X_{i,d}^\nu(t)}{K_d^\nu + X_{i,d}^\nu(t)}\right) \tag{32}$$

Hereby, $\lambda_{A\to B}$ defines the baseline migration rate from tissue $A$ to tissue $B$, with promoting and inhibiting receptor effects following Hill functions, with $K_m$ and $K_d$ defining the half-saturation constants, and the Hill coefficient $\nu$

controlling the sensitivity of the migration rate to changes in marker expression levels. For simplicity, we assume that $\nu$ is identical for all marker and migration processes. The individual migration rates between the different tissues, including the blood, LN, BM, spleen and gut (Suppl. Fig. S1a), with the tissue-specific receptor-ligand interactions as described above are then defined as follows:

$$m_i^{\text{Blood}\to\text{LN}} = \lambda_{\text{Blood}\to\text{LN}} \frac{\text{CCR7}_i^\nu(t)}{K^\nu_{\text{CCR7}} + \text{CCR7}_i^\nu(t)} \frac{\text{CD62L}_i^\nu(t)}{K^n_{\text{CD62L}} + \text{CD62L}_i^\nu(t)} \frac{K^\nu_{\text{S1PR1}}}{K^\nu_{\text{S1PR1}} + \text{S1PR1}_i^\nu(t)} \tag{33}$$

$$m_i^{\text{Blood}\to\text{Spleen}} = \lambda_{\text{Blood}\to\text{Spleen}} \frac{\text{CCR7}_i^\nu(t)}{K^\nu_{\text{CCR7}} + \text{CCR7}_i^\nu(t)} \frac{K^\nu_{\text{S1PR1}}}{K^\nu_{\text{S1PR1}} + \text{S1PR1}_i^\nu(t)} \tag{34}$$

$$m_i^{\text{Blood}\to\text{BM}} = \lambda_{\text{Blood}\to\text{BM}} \frac{\text{CXCR4}_i^\nu(t)}{K^\nu_{\text{CXCR4}} + \text{CXCR4}_i^\nu(t)} \frac{\text{CD44}_i^\nu(t)}{K^\nu_{\text{CD44}} + \text{CD44}_i^\nu(t)} \frac{K^\nu_{\text{S1PR1}}}{K^\nu_{\text{S1PR1}} + \text{S1PR1}_i^\nu(t)} \tag{35}$$

$$m_i^{\text{Blood}\to\text{Gut}} = \lambda_{\text{Blood}\to\text{Gut}} \frac{\alpha4\beta7_i^\nu(t)}{K^\nu_{\alpha4\beta7} + \alpha4\beta7_i^\nu(t)} \frac{K^\nu_{\text{S1PR1}}}{K^\nu_{\text{S1PR1}} + \text{S1PR1}_i^\nu(t)} \tag{36}$$

$$m_i^{\text{LN}\to\text{Blood}} = \lambda_{\text{LN}\to\text{Blood}} \frac{L^\nu_{\text{CCR7}}}{L^\nu_{\text{CCR7}} + \text{CCR7}_i(t)^\nu} \frac{\text{S1PR1}_i^\nu(t)}{L^\nu_{\text{S1PR1}} + \text{S1PR1}_i^\nu(t)} \tag{37}$$

$$m_i^{\text{Spleen}\to\text{Blood}} = \lambda_{\text{Spleen}\to\text{Blood}} \frac{\text{S1PR1}_i^\nu(t)}{L^\nu_{\text{S1PR1}} + \text{S1PR1}_i^\nu(t)} \tag{38}$$

$$m_i^{\text{BM}\to\text{Blood}} = \lambda_{\text{BM}\to\text{Blood}} \frac{L^\nu_{\text{CXCR4}}}{L^\nu_{\text{CXCR4}} + \text{CXCR4}_i^\nu(t)} \frac{L^\nu_{\text{CD44}}}{L^\nu_{\text{CD44}} + \text{CD44}_i^\nu(t)} \frac{\text{S1PR1}_i^\nu(t)}{L^\nu_{\text{S1PR1}} + \text{S1PR1}_i^\nu(t)} \tag{39}$$

$$m_i^{\text{Gut}\to\text{Blood}} = \lambda_{\text{Gut}\to\text{Blood}} \frac{\text{S1PR1}_i^\nu(t)}{L^\nu_{\text{S1PR1}} + \text{S1PR1}_i^\nu(t)} \tag{40}$$

The individual parameters used for the simulations are provided within Table S4, with parameters chosen to approximately match observed tissue dwell times and cell distributions as estimated in [13].

**Model implementation:** The individual processes of marker expression dynamics and cell trafficking between tissues are then simulated using $\tau$-leaping to allow for computationally efficient simulations. For each cell, marker expression levels are updated according to a Poisson distributed random variable representing the number of up- and down-regulation reactions that occur within a fixed time step $\tau$, thus,

$$X_i(t+\tau) = X_i(t) + \kappa - \theta \text{ with } \kappa \sim \text{Pois}(k_{i,x}\tau),\ \theta \sim \text{Pois}(\gamma_{i,x}(l_i|t)X_i(t)\tau) \tag{41}$$

Simultaneously, a cell will transition from tissue $A$ to $B$ at the end of the time interval $\tau$ with probability

$$p_i^{A\to B} = 1 - e^{-m_i^{A\to B}\tau} \tag{42}$$

with the transition event occurring according to randomly sampling from a binomial distribution with Binom($p_i^{A\to B}$). In the case of multiple possible trafficking directions from the originating tissue $A$ (e.g. migration from blood to either lymph node or peripheral tissue), the actual target tissue is sampled according to a multinomial distribution with Eq. (42) updated to

$$p_i^{A\to B} = (1 - p_i(A)) \frac{m_i^{A\to B}}{\sum_B m_i^{A\to B}} \tag{43}$$

with $p_i(A) = e^{-\sum_B m_i^{A\to B}\tau}$ defining the probability of the cell to stay within $A$.

The model was implemented in Python (3.12) with the code available at https://github.com/GrawLab/CellVADIS_analysis.

**Model extensions:** To examine the influence of unobserved tissue sites on cell trafficking fate predictions, we extended the mathematical model introduced above by an additional tissue $U$, which represents the unobserved tissue sites. Migration of cells to this tissue is regulated by a unique marker, $\Lambda$, which also impairs the migration of cells into any of the other sampled tissues, thereby representing the distinct marker profiles between sampled and unobserved tissues. This antagonistic behaviour is included in our model by adapting all migration rates $m_i^{\text{Blood}\to l}$ with $l \neq U$ according to

$$m_i^{\text{Blood}\to l} = m_i^{\text{Blood}\to l} \frac{1}{1 + e^{\gamma(\Lambda_i^{\nu}(t) - K_\Lambda)}} \tag{44}$$

with $\Lambda_i(t)$ denoting the unique marker expression of cell $i$ at time $t$, and $\gamma$, $\nu$ and $K_\Lambda$ defining the sigmoidal function regulating the influence of the marker expression profile. Hereby, the parameter $\gamma$ controls the distinctness of the disparate marker profiles that distinguish cells from the unobserved to all other tissues (see Fig. 3c, Suppl. Fig. S3). Analogously, the migration of blood cells to the unobserved tissue $U$ is described by:

$$m_i^{\text{Blood}\to U} = \lambda_{\text{Blood}\to U} \frac{1}{1 + e^{-\gamma(\Lambda_i^{\nu}(t) - K_\Lambda)}} \frac{K_{\text{S1PR1}}^{\nu}}{K_{\text{S1PR1}}^{\nu} + \text{S1PR1}_i^{\nu}(t)}, \tag{45}$$

with cells leaving tissue $U$ again with the default exit rate

$$m_i^{U\to\text{Blood}} = \lambda_{U\to\text{Blood}} \frac{\text{S1PR1}_i^{\nu}(t)}{L_{\text{S1PR1}}^{\nu} + \text{S1PR1}_i^{\nu}(t)}. \tag{46}$$

**Table S1: Model hyperparameters:** General parameters for simulating individual cell-based marker dynamics

| Parameter | Value | Prior | Unit | Description |
|---|---|---|---|---|
| $N_{\text{total}}$ | 50000 | $\mathcal{U}(10^4, 10^5)$ | – | Number of cells |
| $\tau^{\text{leap}}$ | 0.02 | $\mathcal{U}(0.01, 0.1)$ | h | Time step for tau-leaping algorithm |
| $\nu$ | 3 | $\mathcal{U}(1, 6)$ | – | Hill coefficient for all markers and migration processes |

**Table S2: Marker-specific expression and degradation rates:** For each of the markers considered, the parameters for describing marker expression and degradation as used in the corresponding expressions of Eq. (23) are given. (*KC: half-saturation constant*).

| Marker | Parameter | Value | Prior | Unit | Description |
|---|---|---|---|---|---|
| CD44 | $\rho_{0,\text{CD44}}$ | 5 | – | $h^{-1}$ | Base production rate |
| | $\delta_{0,\text{CD44}}$ | 0.1 | – | $h^{-1}$ | Degradation rate |
| CCR7 | $\rho_{0,\text{CCR7}}$ | 8 | – | $h^{-1}$ | Base production rate |
| | $\delta_{0,\text{CCR7}}$ | 0.1 | – | $h^{-1}$ | Basal degradation rate |
| | $\delta_{1,\text{CCR7}}$ | 1 | – | $h^{-1}$ | Internalization rate in lymphoid organs |
| | $T^{\nu}_{\text{CCR7}}$ | 8 | $\mathcal{U}(2, 16)$ | h | Time constant for internalization dynamics |
| S1PR1 | $\rho_{0,\text{S1PR1}}$ | 12 | – | $h^{-1}$ | Base production rate |
| | $\delta_{0,\text{S1PR1}}$ | 0.1 | – | $h^{-1}$ | Basal degradation rate |
| | $\delta_{1,\text{S1PR1}}$ | 1 | – | $h^{-1}$ | Internalization rate in blood |
| | $T^{\nu}_{\text{S1PR1}}$ | 0.05 | $\mathcal{U}(0.01, 1)$ | h | Time constant for internalization in blood |
| | $K^{\nu}_{\text{CD69}}$ | 60 | – | – | KC for CD69-mediated degradation |
| | $\nu_{\text{CD69}}$ | 6 | – | – | Hill coefficient for CD69-mediated degradation |
| CXCR4 | $\rho_{0,\text{CXCR4}}$ | 6 | – | $h^{-1}$ | Base production rate |
| | $\delta_{0,\text{CXCR4}}$ | 0.1 | – | $h^{-1}$ | Basal degradation rate |
| | $\delta_{1,\text{CXCR4}}$ | 0.5 | – | $h^{-1}$ | Additional degradation in bone marrow |
| | $T^{\nu}_{\text{CXCR4}}$ | 16 | $\mathcal{U}(2, 16)$ | h | Time constant for internalization dynamics |
| $\alpha 4\beta 7$ | $\rho_{0,\alpha 4\beta 7}$ | 4 | – | $h^{-1}$ | Base production rate |
| | $\delta_{0,\alpha 4\beta 7}$ | 0.1 | – | $h^{-1}$ | Degradation rate |
| CD69 | $\rho_{0,\text{CD69}}$ | 3 | – | $h^{-1}$ | Base production rate |
| | $\delta_{0,\text{CD69}}$ | 0.1 | – | $h^{-1}$ | Degradation rate |
| CD62L | $\rho_{0,\text{CD62L}}$ | 8 | – | $h^{-1}$ | Base production rate |
| | $\delta_{0,\text{CD62L}}$ | 0.1 | – | $h^{-1}$ | Degradation rate |
| $\Lambda$ | $\rho_{0,\Lambda}$ | 4 | – | $h^{-1}$ | Base production rate |
| | $\delta_{0,\Lambda}$ | 0.1 | – | $h^{-1}$ | Degradation rate |

**Table S3: Parameters accounting for cell-to-cell variability in marker expression:** Expression multipliers for each marker are sampled from a bimodal distribution consisting generally of a Lognormal- and a Normal-distributed part, with exception for S1PR1 only being sampled from a Lognormal-distribution.

| Marker | Parameter | Value | Prior | Unit | Description |
|---|---|---|---|---|---|
| CD44 | $\lambda^{\mathrm{CD44}}$ | 0.70 | $\mathcal{U}(0.1, 1)$ | – | Weight for Lognormal component (low expression) |
| | $\mu_{\mathrm{G}}^{\mathrm{CD44}}$ | 2.0 | – | – | Mean of Gaussian component |
| | $\sigma_{\mathrm{G}}^{\mathrm{CD44}}$ | 0.3 | – | – | SD of Gaussian component |
| | $\mu_{\mathrm{L}}^{\mathrm{CD44}}$ | -1.0 | – | – | Mean of Lognormal component |
| | $\sigma_{\mathrm{L}}^{\mathrm{CD44}}$ | 0.5 | – | – | SD of Lognormal component |
| CCR7 | $\lambda^{\mathrm{CCR7}}$ | 0.30 | $\mathcal{U}(0.1, 1)$ | – | Weight for Lognormal component |
| | $\mu_{\mathrm{G}}^{\mathrm{CCR7}}$ | 1.7 | – | – | Mean of Gaussian component |
| | $\sigma_{\mathrm{G}}^{\mathrm{CCR7}}$ | 0.3 | – | – | SD of Gaussian component |
| | $\mu_{\mathrm{L}}^{\mathrm{CCR7}}$ | -3.0 | – | – | Mean of Lognormal component |
| | $\sigma_{\mathrm{L}}^{\mathrm{CCR7}}$ | 0.5 | – | – | SD of Lognormal component |
| S1PR1 | $\mu_{\mathrm{L}}^{\mathrm{S1PR1}}$ | 0 | – | – | Mean of Lognormal distribution |
| | $\sigma_{\mathrm{L}}^{\mathrm{S1PR1}}$ | 0.1 | – | – | SD of Lognormal distribution |
| CXCR4 | $\lambda^{\mathrm{CXCR4}}$ | 0.20 | $\mathcal{U}(0.1, 1)$ | – | Weight for Lognormal component |
| | $\mu_{\mathrm{G}}^{\mathrm{CXCR4}}$ | 1.8 | – | – | Mean of Gaussian component |
| | $\sigma_{\mathrm{G}}^{\mathrm{CXCR4}}$ | 0.8 | – | – | SD of Gaussian component |
| | $\mu_{\mathrm{L}}^{\mathrm{CXCR4}}$ | -1.0 | – | – | Mean of Lognormal component |
| | $\sigma_{\mathrm{L}}^{\mathrm{CXCR4}}$ | 0.5 | – | – | SD of Lognormal component |
| CD62L | $\lambda^{\mathrm{CD62L}}$ | 0.30 | $\mathcal{U}(0.1, 1)$ | – | Weight for Lognormal component |
| | $\mu_{\mathrm{G}}^{\mathrm{CD62L}}$ | 1.9 | – | – | Mean of Gaussian component |
| | $\sigma_{\mathrm{G}}^{\mathrm{CD62L}}$ | 0.4 | – | – | SD of Gaussian component |
| | $\mu_{\mathrm{L}}^{\mathrm{CD62L}}$ | -1.0 | – | – | Mean of Lognormal component |
| | $\sigma_{\mathrm{L}}^{\mathrm{CD62L}}$ | 0.6 | – | – | SD of Lognormal component |
| $\alpha 4\beta 7$ | $\lambda^{\alpha 4\beta 7}$ | 0.50 | $\mathcal{U}(0.1, 1)$ | – | Weight for Lognormal component |
| | $\mu_{\mathrm{G}}^{\alpha 4\beta 7}$ | 1.9 | – | – | Mean of Gaussian component |
| | $\sigma_{\mathrm{G}}^{\alpha 4\beta 7}$ | 0.4 | – | – | SD of Gaussian component |
| | $\mu_{\mathrm{L}}^{\alpha 4\beta 7}$ | -1.0 | – | – | Mean of Lognormal component |
| | $\sigma_{\mathrm{L}}^{\alpha 4\beta 7}$ | 0.6 | – | – | SD of Lognormal component |
| CD69 | $\lambda^{\mathrm{CD69}}$ | 0.50 | $\mathcal{U}(0.1, 1)$ | – | Weight for Lognormal component |
| | $\mu_{\mathrm{G}}^{\mathrm{CD69}}$ | 1.9 | – | – | Mean of Gaussian component |
| | $\sigma_{\mathrm{G}}^{\mathrm{CD69}}$ | 0.4 | – | – | SD of Gaussian component |
| | $\mu_{\mathrm{L}}^{\mathrm{CD69}}$ | -1.0 | – | – | Mean of Lognormal component |
| | $\sigma_{\mathrm{L}}^{\mathrm{CD69}}$ | 0.6 | – | – | SD of Lognormal component |
| $\Lambda$ | $\lambda^{\Lambda}$ | 0.80 | $\mathcal{U}(0.1, 1)$ | – | Weight for Lognormal component |
| | $\mu_{\mathrm{G}}^{\Lambda}$ | 2.1 | – | – | Mean of Gaussian component |
| | $\sigma_{\mathrm{G}}^{\Lambda}$ | 0.4 | – | – | SD of Gaussian component |
| | $\mu_{\mathrm{L}}^{\Lambda}$ | -1.0 | – | – | Mean of Lognormal component |
| | $\sigma_{\mathrm{L}}^{\Lambda}$ | 0.6 | – | – | SD of Lognormal component |

**Table S4: Parameters describing tissue-specific migration dynamics:** Model parameters used to simulate tissue-specific migration dynamics as described in Eq. (33) - (40). (EC: half-saturation constant).

| Parameter | Value | Prior | Unit | Description |
|---|---|---|---|---|
| General parameters | | | | |
| $K_{\mathrm{S1PR1}}$ | 40 | – | – | EC for S1PR1 repression (inhibitory) (all reactions *Blood* →) |
| $L_{\mathrm{S1PR1}}$ | 60 | – | – | EC for S1PR1-mediated egress (all reactions → *Blood*) |
| Blood → Lymph Node | | | | |
| $\lambda_{\mathrm{Blood}\to\mathrm{LN}}$ | 30 | $\mathcal{U}(1, 100)$ | $h^{-1}$ | Baseline migration rate |
| $K_{\mathrm{CCR7}}$ | 100 | $\mathcal{U}(40, 160)$ | – | EC for CCR7-mediated migration |
| $K_{\mathrm{CD62L}}$ | 130 | $\mathcal{U}(40, 160)$ | – | EC for CD62L-mediated migration |
| Lymph Node → Blood | | | | |
| $\lambda_{\mathrm{LN}\to\mathrm{Blood}}$ | 0.07 | $\mathcal{U}(0.01, 1)$ | $h^{-1}$ | Baseline egress rate |
| $L_{\mathrm{CCR7}}$ | 70 | – | – | EC for CCR7 repression (inhibitory) |
| Blood → Spleen | | | | |
| $\lambda_{\mathrm{Blood}\to\mathrm{Spleen}}$ | 15 | $\mathcal{U}(1, 100)$ | $h^{-1}$ | Baseline migration rate |
| $K_{\mathrm{CCR7}}$ | 100 | $\mathcal{U}(40, 160)$ | – | EC for CCR7-mediated migration |
| Spleen → Blood | | | | |
| $\lambda_{\mathrm{Spleen}\to\mathrm{Blood}}$ | 1.5 | $\mathcal{U}(0.01, 2)$ | $h^{-1}$ | Baseline egress rate |
| Blood → Bone Marrow | | | | |
| $\lambda_{\mathrm{Blood}\to\mathrm{BM}}$ | 9 | $\mathcal{U}(1, 100)$ | $h^{-1}$ | Baseline migration rate |
| $K_{\mathrm{CXCR4}}$ | 120 | $\mathcal{U}(40, 160)$ | – | EC for CXCR4-mediated migration |
| $K_{\mathrm{CD44}}$ | 90 | $\mathcal{U}(40, 160)$ | – | EC for CD44-mediated migration |
| Bone Marrow → Blood | | | | |
| $\lambda_{\mathrm{BM}\to\mathrm{Blood}}$ | 0.04 | $\mathcal{U}(0.01, 1)$ | $h^{-1}$ | Baseline egress rate |
| $L_{\mathrm{CXCR4}}$ | 120 | $\mathcal{U}(40, 160)$ | – | EC for CXCR4 repression (inhibitory) |
| $L_{\mathrm{CD44}}$ | 100 | $\mathcal{U}(40, 160)$ | – | EC for CD44 repression (inhibitory) |
| Blood → Gut | | | | |
| $\lambda_{\mathrm{Blood}\to\mathrm{Gut}}$ | 6 | $\mathcal{U}(1, 100)$ | $h^{-1}$ | Baseline migration rate |
| $K_{\alpha 4\beta 7}$ | 80 | $\mathcal{U}(40, 160)$ | – | EC for $\alpha4\beta7$-mediated migration |
| Gut → Blood | | | | |
| $\lambda_{\mathrm{Gut}\to\mathrm{Blood}}$ | 0.2 | $\mathcal{U}(0.01, 1)$ | $h^{-1}$ | Baseline egress rate |
| Blood → U | | | | |
| $\lambda_{\mathrm{Blood}\to\mathrm{U}}$ | 6 | $\mathcal{U}(1, 100)$ | $h^{-1}$ | Baseline migration rate |
| $\lambda$ | 1 | $\mathcal{U}(10^{-3}, 1)$ | – | Distinctness parameter |
| $K_{\Lambda}$ | 50 | | – | Value where sigmoidal term is 0.5 |
| U → Blood | | | | |
| $\lambda_{\mathrm{U}\to\mathrm{Blood}}$ | 0.2 | $\mathcal{U}(0.01, 1)$ | $h^{-1}$ | Baseline egress rate |

#### Text S2: Impact of unobserved tissue sites on cell fate prediction

As outlined in the main manuscript, cell fate prediction for cells that would actually migrate to sampled tissues would not be affected by unobserved tissues sites. This observation is not surprising for *StitchOT*, as here, by definition location fates are inferred independently from each other due to the tissue-specific OT-maps. Therefore, adding an additional column for the unobserved tissue $U$ to the fate matrix only downscales the values to all other sampled tissues, i.e.

$$p_i^{\mathrm{wU}}(l) = (1 - p_i^{\mathrm{wU}}(U))p_i^{\mathrm{w/oU}}(l) \tag{47}$$

For the probabilities conditioned on not going to $U$ this implies

$$p_i^{\mathrm{wU}}(l|\neg U) = \frac{p^{\mathrm{wU}}(l \cap \neg U)}{p_i^{\mathrm{wU}}(\neg U)} = \frac{p_i^{\mathrm{wU}}(l)}{1 - p_i^{\mathrm{wU}}(U)} = p_i^{\mathrm{w/oU}}(l) \tag{48}$$

Here $p_i^{\mathrm{wU}}(l)$ denotes the fate probability for cell $i$ to transition to location $l \in \{1, \ldots, L\}$ given full information, i.e. considering samples from the unobserved tissue $U$. Analogously, $p_i^{\mathrm{w/oU}}$ refers to the inferred fate probability without accounting for cells from $U$, i.e. not considering the unobserved tissue.

Although there is no such theoretical guarantee for *JointOT*, robustness tests in Fig. 3d show that the conditional probabilities $p_i^{\mathrm{wU}}$ and $p_i^{\mathrm{w/oU}}$ do not differ significantly, indicating that the incorrect mass distribution caused by mis-mapped cells ($l_i^{\mathrm{next}} = U$) does not alter the fate assignment for the other cells ($l_i^{\mathrm{next}} \neq U$).

### Text S3: OT-method variations to assess method robustness

In order to assess the robustness and appropriateness of our presented method formulations and parameterisations of *StitchOT* and *JointOT*, we tested several variations of the chosen frameworks and compared their performance in the ability to predict cell fates.

This included the following variations:

- **Marginal constraints:** Varying the marginal constraints $\tau_a$ and $\tau_b$ for *StitchOT* and *JointOT*, respectively, with increasing values converging towards balanced optimal transport methods not allowing for mass loss or gain between the blood and tissue samples. This clearly leads to a decrease in model performance (Suppl. Fig. S2), indicating that the unbalanced formulation of OT is necessary to correctly map cells across tissues, i.e. accounting for cell proliferation and differentiation within individual tissues, as well as biased sampling.
- **Rescaling:** Testing the benefit of using estimated weighting factors $\omega_l$ (Eq. (6)) in *StitchOT* for the location-specific OT-maps to rescale the global OT-map $T$, by neglecting rescaling, i.e. $\omega_l = 1$ for $l \in \{1, \ldots, L\}$.
- **Subsampling:** A further aspect considered in our methods was the subsampling of cells from the different tissues in order to ensure equal sample sizes for avoiding potential bias due to unbalanced tissue sizes. Although this is accounted for in our methods by rescaling the marginal weights such that $b_i = \frac{1}{L}\frac{1}{M_{l_i}}$, the entropy term enforcing the marginals, depends on the scale of the marginal values. Therefore, providing this kind of prior knowledge could influence weight distributions in favor of tissues with higher cell counts.
- **Backward-Mapping:** Due to having no temporal information on which cells are about to enter or leave a particular tissue, CellVADIS currently could map cells that are close to leaving the blood 'backwards' to cells that are about to leave the tissue instead of to tissue cells that recently entered. To test the extent to which this kind of backward mapping could impair fate prediction, we filtered cells based on their S1PR1 expression such that only source cells that are about to enter a tissue and target cells that recently entered a tissue according to the interactions implemented in our model, i.e. only considering cells below the 33-rd percentile of S1PR1-expression within the analysis.
- **Tissue proportions:** If prior knowledge about the proportion of blood cells that would migrate to a specific tissue would be available, this information could be used to define the tissue-specific weights, $\omega_l$, or marginal weights in *StitchOT* or *JointOT*, respectively, which could potentially improve fate predictions due to better tissue mass distribution.

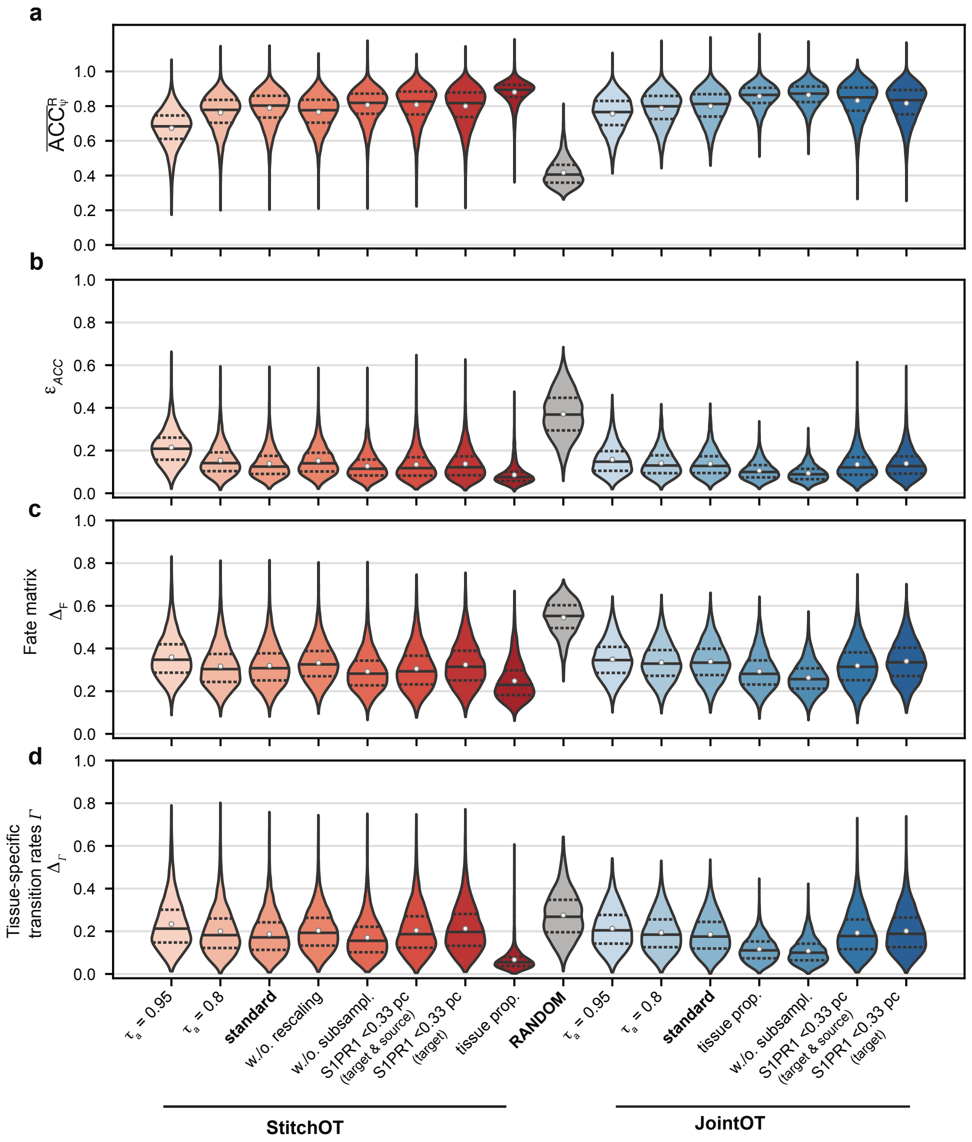


**Figure S2: Accuracy of cell fate prediction according to variations in the methodological framework:** Assessed cell fate prediction ability for *StitchOT* and *JointOT* in comparison to variations in the methodological framework addressing selection of marginal constraints, scaling, cell subsampling, backward-mapping and information on tissue proportions (see Suppl. Information Text S3). Performance measures including the relative average threshold accuracy, $\overline{\text{ACC}}^{\text{R}}_{\psi}$ (**a**), and integrated absolute error, $\varepsilon_{\text{ACC}}$ (**b**), as well as the total variation in transition probabilities, $\Delta_{\text{F}}$ (**c**), and tissue-specific transition proportions, $\Delta_{\Gamma}$ (**d**), are shown in correspondence to Fig. 2f.

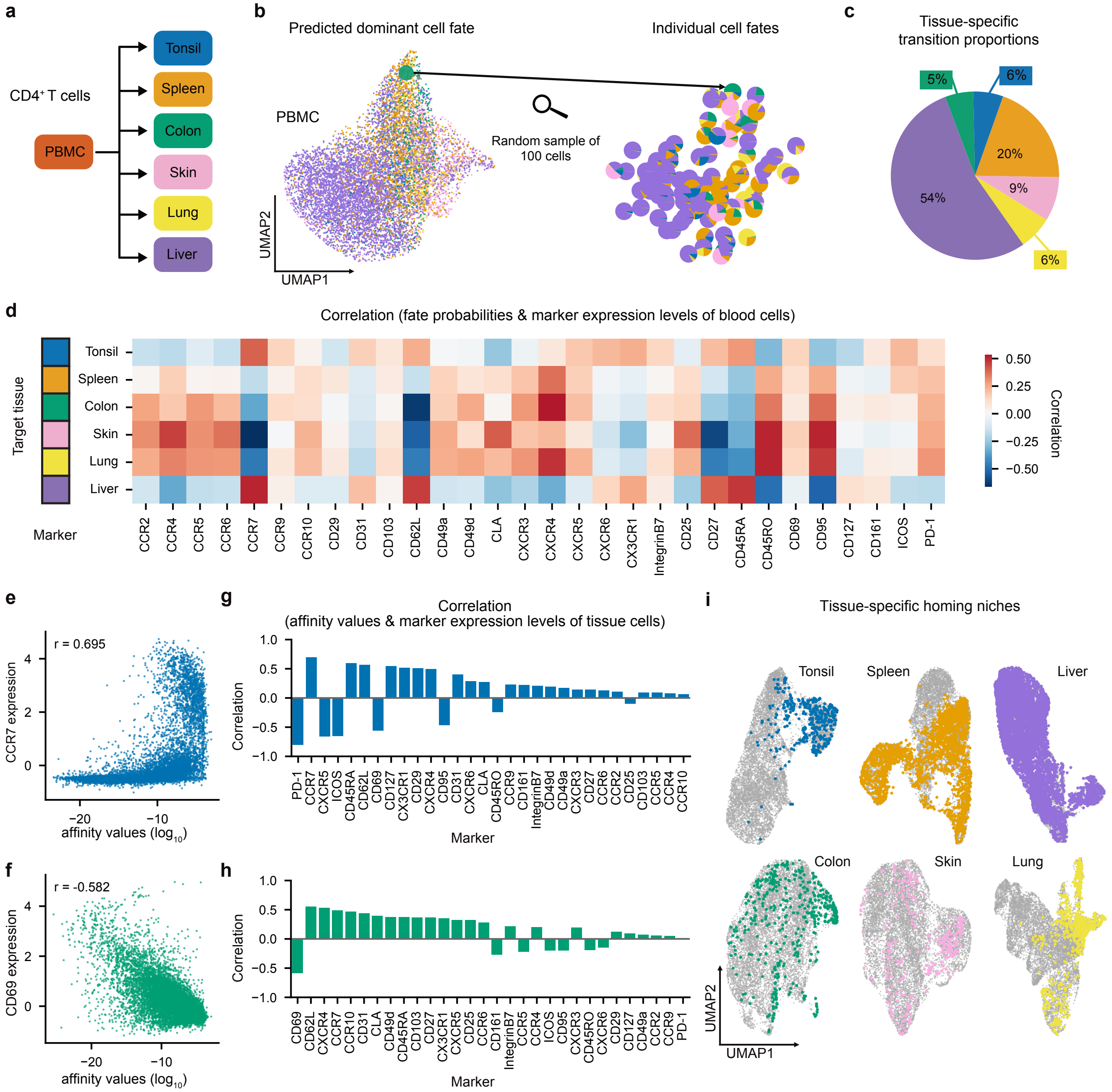


**Figure S4: Cell fate prediction based on mass cytometry data for CD4$^+$ T cells (Wong et al. 2016):** (**a**) Tissue sites considered by Wong et al. [14] with PBMC measurements (=blood cells) used for fate prediction. (**b**) UMAP representation of $10^4$ CD4$^+$ T cells of the PBMC coloured by the dominant fate prediction by *JointOT* with individual fate predictions for 100 randomly sampled cells. (**c**) Predicted tissue-specific transition proportions for PBMC measurements. (**d**) Heatmap of the correlation of specific marker expression levels and predicted cell fates for the different tissues. (**e-f**) Exemplary correlation of marker expression levels for CCR7 and CD69 of individual tissue CD4$^+$ T cells for the tonsil (**e**) and colon (**f**), respectively, with their affinity values towards blood cells. (**g-h**) Corresponding correlation values across the tissue-specific markers for tonsil (**g**) and colon (**h**). (**i**) Identification of homing niches of PBMC-CD4$^+$ T cells with a high fate potential ($\psi \geq 1/3$) to the particular tissue by projection into the UMAP representation of the CD4$^+$ T cells measured at the respective tissue site.

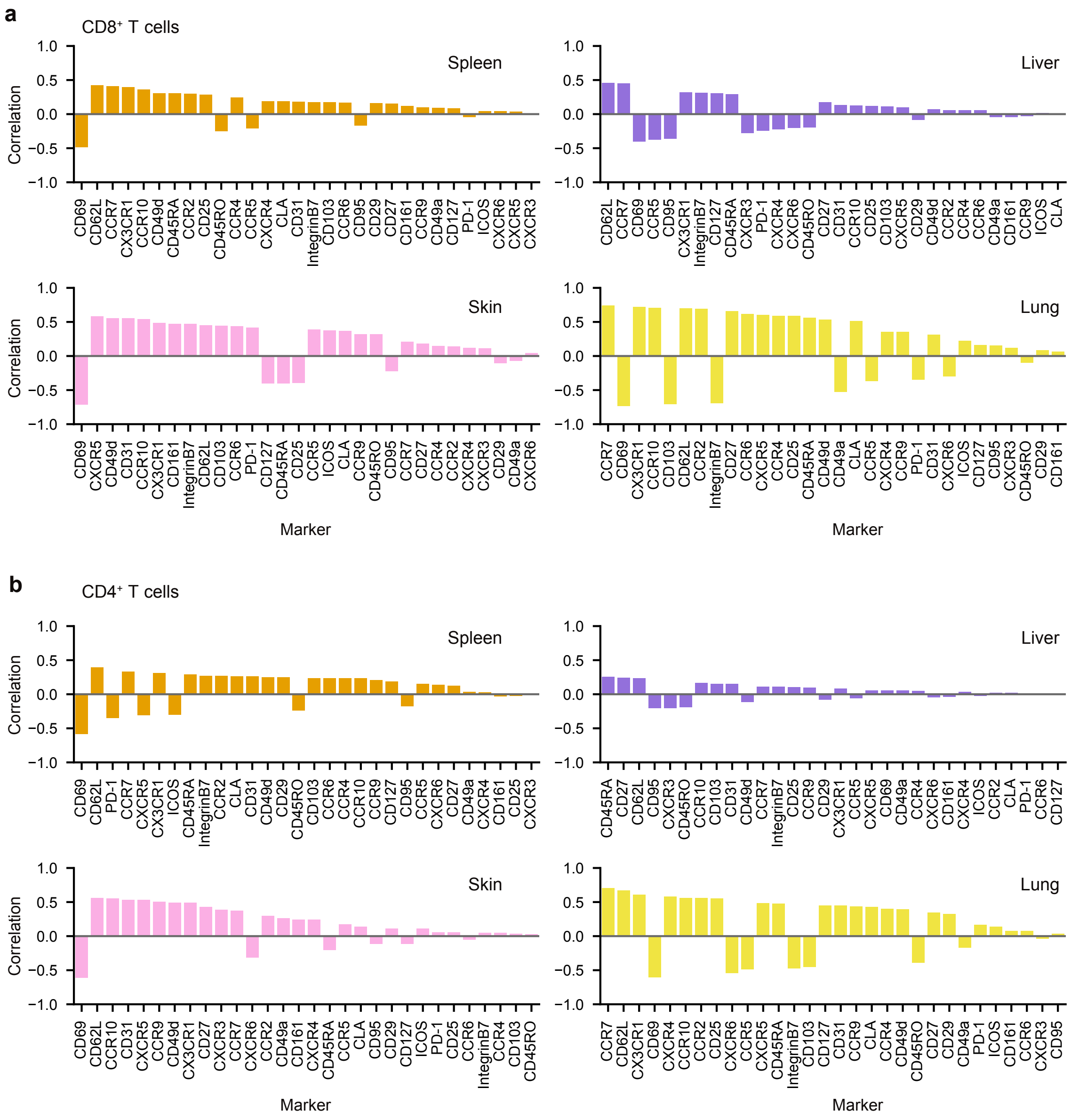


**Figure S5: Trafficking signatures for different tissues and cell types:** Correlation between affinity values and marker expression levels for tissue $CD8^+$ (**a**) and $CD4^+$ (**b**) T cells for the additional tissues not shown in Fig. 4g,h and Suppl. Fig. S4, respectively.

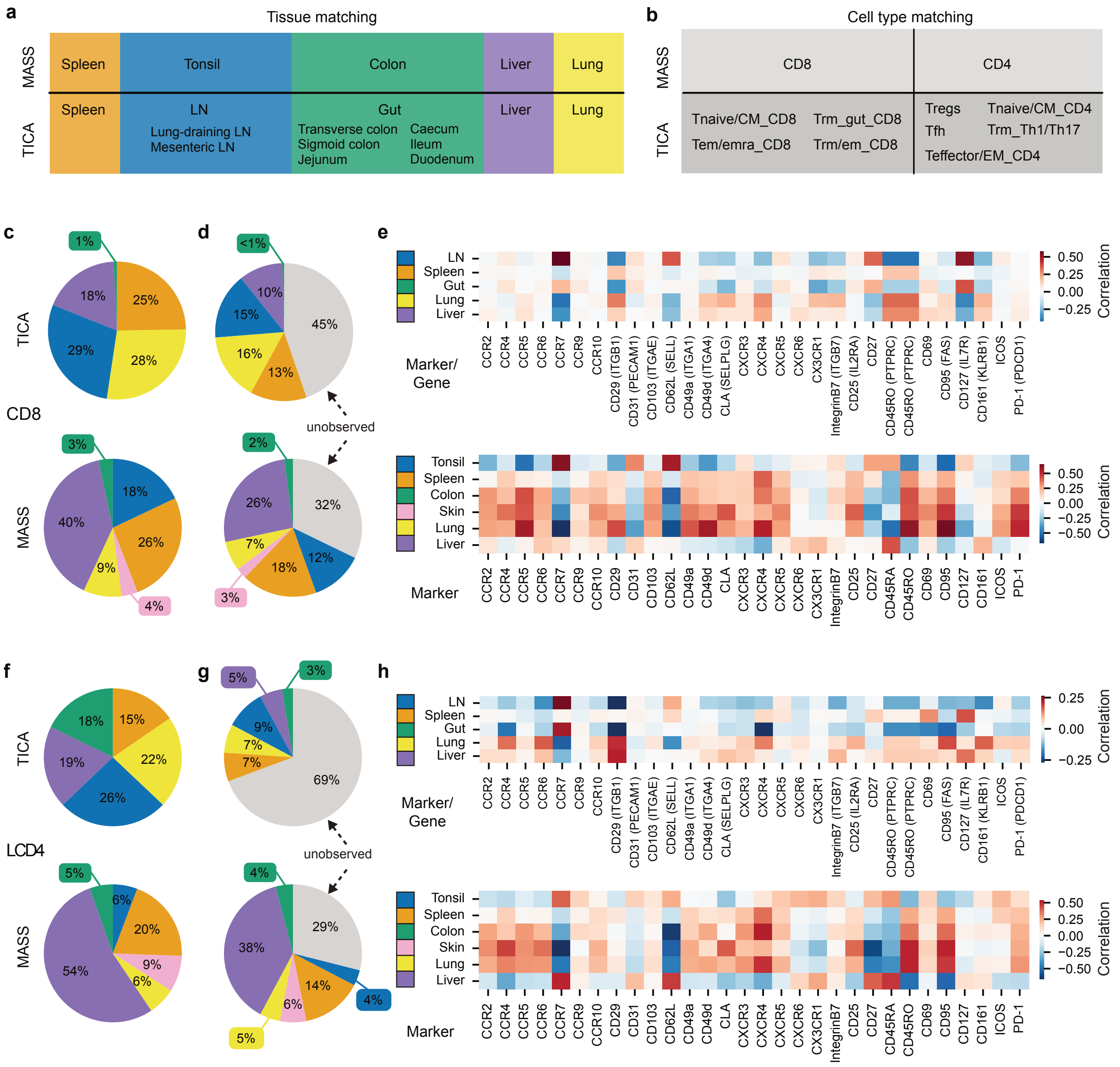


**Figure S6: Comparing cell fate prediction based on different data types:** (**a**) Alignment of different tissues given for mass cytometry analyses (MASS) [14] and the Cross-Tissue Immune cell Atlas (TICA) based on scRNAseq-measurements[15]. (**b**) Combination of manually curated cell types for comparison of both datasets. (**c-d**) Predicted tissue-specific transition proportions for CD8$^+$ T cells with (**c**) and without (**d**) assuming the occurrence of unobserved tissue sites applying *StitchOT*. (**e**) Correlation of tissue-specific cell fate predictions with marker (MASS) or corresponding gene expression levels (TICA). (**f-h**) Analogous analyses for CD4$^+$ T cells corresponding to panel (**c-e**), respectively.

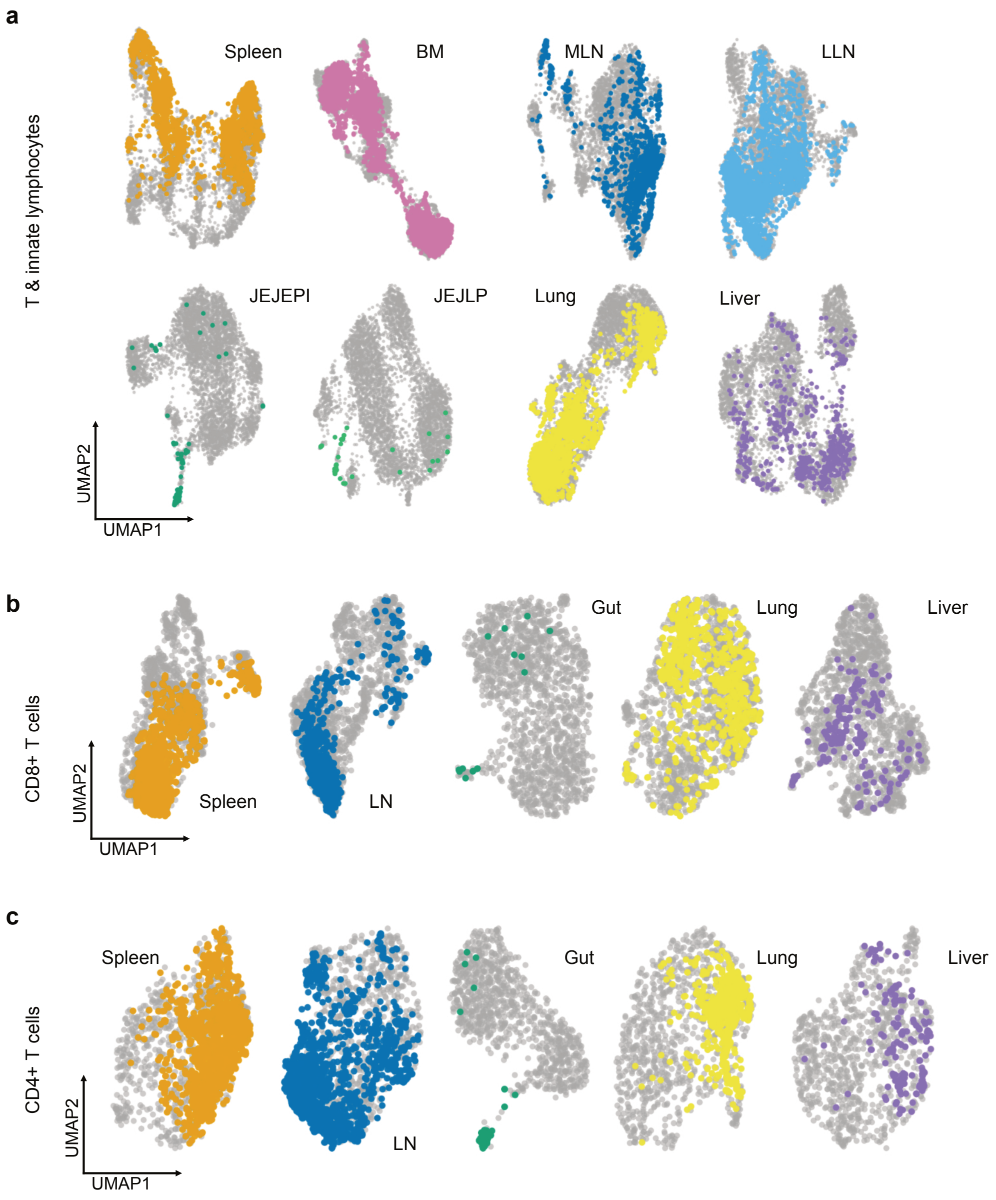


**Figure S7: Prediction of homing niches for T & innate lymphocytes based on the cross-tissue immune atlas (TICA):** UMAP-projection of blood cells into the corresponding tissues according to their fate predictions considering all T & innate lymphocytes (**a**), as well as restricted to $CD8^+$ T cell **(b)** and $CD4^+$ T cells (**c**) [15]. See Suppl. Fig. S6 for details on chosen subsets and tissue alignment.
